# Impact of Aii Amacrine Cell Rewiring in a Pathoconnectome-Based Computational Model of Early Retinal Degeneration

**DOI:** 10.1101/2024.02.14.580149

**Authors:** Ege Iseri, Rebecca Pfeiffer, Crystal Sigulinsky, James Anderson, Jia-Hui Yang, Jeebika Dahal, Jessica Garcia, Jean-Marie C. Bouteiller, Bryan W. Jones, Gianluca Lazzi

## Abstract

Retinitis pigmentosa (RP), a retinal degenerative disease, is characterized by progressive photoreceptor loss and ongoing remodeling and rewiring of the inner retina. This study investigates rod network rewiring through pathoconnectomic evaluation and its impacts on signaling patterns. The glycinergic Aii amacrine cell (Aii) plays a central role in the healthy retina bridging rod and cone pathways, enabling an increased dynamic range of vision. Pathoconnectomics reveals altered connectivity in both the excitatory drive and gap junctional coupling of Aiis in retinal degeneration. A computational model of the rewired network was developed to assess the functional consequences of these structural changes by simulating light-evoked responses and changes in excitatory postsynaptic potentials (EPSPs). The model predicts significant changes in bipolar and Aii EPSPs between active and baseline conditions, driven by newly formed gap junctions in the degenerate retina. Notably, the aberrant circuitry induces rhythmic firing of up to 10 Hz in retinal ganglion cells, consistent with network depolarization relative to the healthy baseline state. These findings align with patch-clamp observations in rd1 and rd10 mouse models of RP, suggesting that Aii-mediated network alterations may underlie early clinical symptoms, including impaired adaptation between photopic and scotopic vision. More broadly, this work demonstrates that integrating computational modeling with pathoconnectomics enables predictive analysis of signaling in early-stage retinal degeneration and may help identify windows for therapeutic intervention. Such models could be further extended with multi-scale bioelectromagnetic simulations to optimize neurostimulation strategies aimed at slowing disease progression.

**Author summary:** Understanding how retinal degeneration alters wiring topologies of the inner retina is important for the success of multiple therapeutic interventions, including cell replacement strategies, optogenetics, and electrode implants. Here, we continue our evaluation of retinal pathoconnectome 1 (RPC1), describing additional network-level changes occurring early in retinal degeneration. This analysis extends our previous findings on the emergence of gap junctions in rod bipolar cells in retinal degeneration to include the effects of these changes on the synaptic strength of inputs to the Aii. We then model how these network changes overall effect retinal processing through the creation of a more complete degenerate retina model. From these results we propose the emergence of aberrant gap junctions in the rod pathway as the network cause of atypical retinal ganglion cell firing and provide the field with a realistic network model for evaluating and optimizing therapeutic strategies.

## Introduction

The retina is a complex, layered structure that converts light into electrical signals through phototransduction. This is followed by signal processing in the neural retina, breaking down visual scenes into its composite parts, thereby enabling visual perception. The retinal neural network is comprised of various cell classes connected through excitatory and inhibitory chemical synapses and electrical synapses. Retinitis Pigmentosa (RP) is a retinal degenerative disease caused by numerous genetic mutations resulting in the progressive loss of rod photoreceptors [1]. Along with the loss of rods, the retina enters into a state of retinal remodeling [2]. There are many components of retinal remodeling, including gliosis, cell death, cell migration, and rewiring, that encompass all retinal classes of neurons and eventually progress to a wide-spread neurodegeneration [3]. Of particular relevance to this study, is the process of rewiring, wherein neurons make aberrant connections in the diseased retina [2, 4–9]. In advanced retinal degeneration, neurons will extend neurite processes into unexpected retinal layers and make synapses with unknown partners [2]. This rewiring is observed in animal models of retinal degeneration with genetic causes [4, 7, 8], as well as induced non-genetic causes [10], and has been confirmed in human RP [11] and AMD patients [12].

Within the retinal network, the glycinergic Aii amacrine cell (Aii) is a central node for visual processing [13]. In healthy mammalian retina, the Aii mediates scotopic (low-light) vision by relaying signals from rod photoreceptor-connected rod bipolar cells (RodBCs) to retinal ganglion cell (RGC)-connected ON-cone bipolar cells (CBbs) by bridging chemical synapses with gap junctions, or electrical synapses [13–15]. The Aii engages in reciprocal synapses with OFF-cone bipolar cells (CBas), contributing to scotopic-driven OFF signaling, ON-OFF opponency, and rod-cone crossover opponency through the lateral inhibition of CBas [16]. Additionally, studies have demonstrated extensive gap junctional coupling between Aiis [16–18], and that gap junctional coupling strength varies with luminance [19, 20].

Given the paucity of direct synapses between RodBCs and RGCs in the healthy mammalian retina, the connection to the CBbs via gap junctions with the Aii is the primary mechanism of light perception under scotopic conditions. When this pathway is disrupted, significant loss of scotopic vision occurs [21, 22] The function of the Aii in ON-OFF and rod-cone crossover motifs is more subtle, yet observable through psychophysical analysis of cone flicker threshold during rod adaptation. As rods reach a dark adaptation state, the intensity of light required for flicker perception via cone input is increased [23], presumably through inhibition of the cone pathway in the inner retina by the rod pathway. Aiis participate in surround inhibition over most of the visual dynamic range, except the most scotopic of conditions [24]. Disruption of this pathway component has been observed in patients with Retinitis Pigmentosa (RP) and other rod dystrophies [25].

Understanding the precise network topologies which emerge in retinal rewiring is important for designing therapeutic interventions that can be used across retinal degenerative diseases. Despite significant progress in retinal prosthetics, there remains substantial room for improvement [26–28]. A major challenge with the current technology is reproducing the natural light response of the retina through electrical stimulation waveforms. Electrode array resolution limits selective stimulation, and using smaller electrodes does not solve this problem [29, 30]. Stimulation parameters such as pulse width, frequency, and phase balance [31–33] must be optimized to reproduce the natural light responses in the target cells. One strategy to improve implant effectiveness is designing waveforms that stimulate local cell clusters and approximate spatiotemporal activation patterns that better represent the input image [34]. Achieving this requires a realistic retinal network model that contains the relevant cell types, accurate membrane properties, and precise connectivity mapping.

Here, we further explored Retinal Pathoconnectome 1: *RPC1* [35] computationally to interrogate the state of rewiring in the Aii networks early in retinal degeneration. A computational retina network model was constructed to investigate how retinal signaling changes during degeneration. The model incorporates realistic morphologies and topologies of the selected cells from RPC1, specifically the aberrant synapses that occur exclusively in degenerate retina. A current injection protocol was developed to capture the retina’s response to light stimulation, using patch-clamp recordings of each cell type under the same conditions. The key component of early-stage retinal degeneration simulated here are the aberrant gap junctions between Aiis and RodBCs due to rewiring [35]. We evaluate the impact of these gap junctions on the network activity and highlight the differences from a healthy retina baseline.

## 1 RPC1 Aii Network Evaluation Results

### 1.1 Gross Anatomy of Aii Cells

RPC1 contains 11 Aiis. To quantify the most accurate synapse numbers and weights per Aii, the 5 Aiis that are predominately contained within the volume (cells minimally have processes extending beyond the bounds of the volume) were reconstructed (Cell #s 192, 262, 265, 1685, 2710). These cells were compared against representative Aiis from our previously published retinal connectome 1: *RC1*. RC1 is a robust connectomics dataset generated from a healthy rabbit retina and has been extensively annotated, providing comprehensive descriptions of Aii connectivity motifs [13, 36]. All of the identified Aiis in the RPC1 volume contain moderate glycine levels, similar to that found within Aiis of *RC1* [13] (Figure 1), and consistent with prior studies [37, 38].

**Fig 1.**
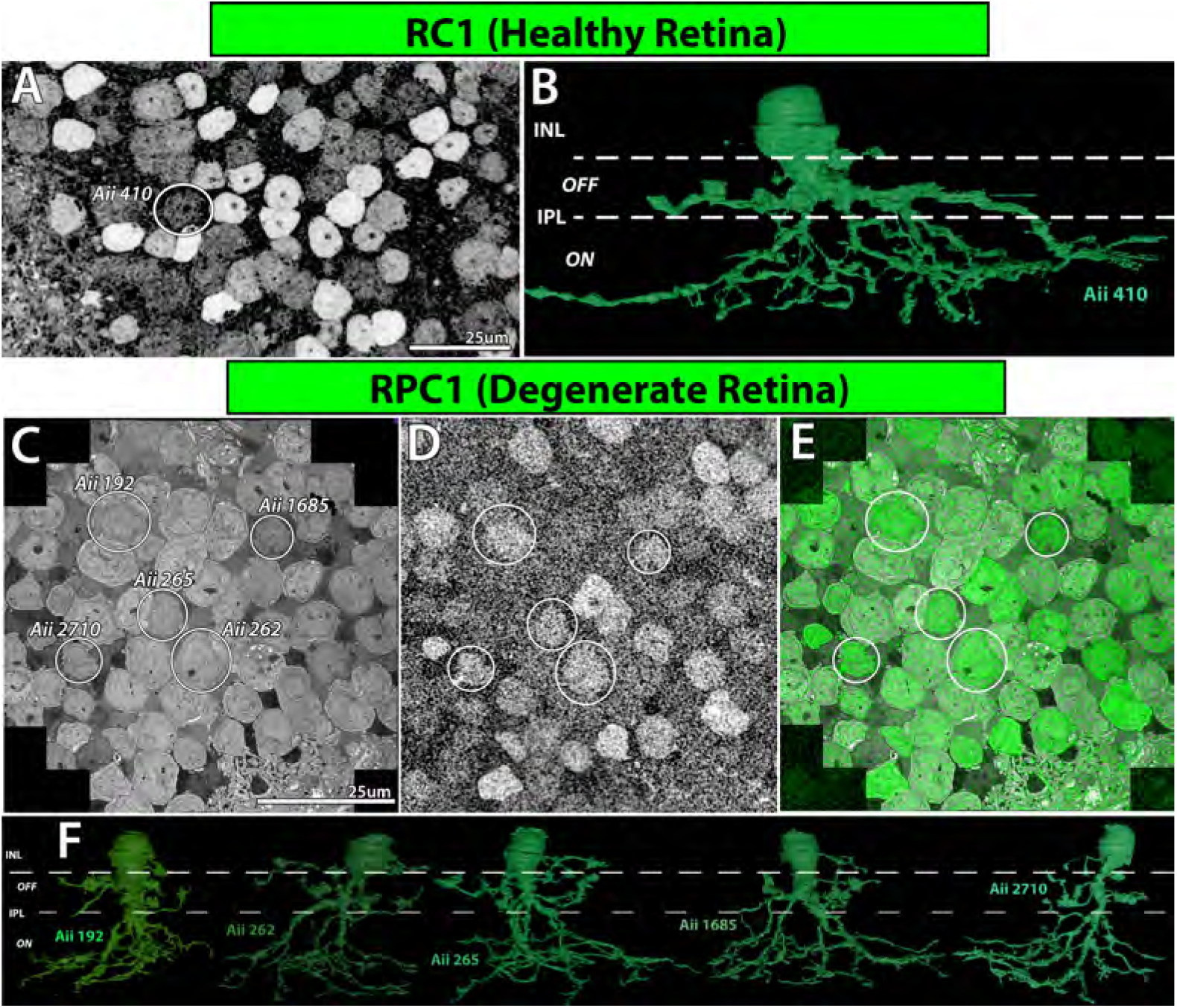
Aii Morphology and Glycine Content. (A) Glycine-labeled section (section 30) demonstrated the medium levels of glycine found in RC1 Aiis, including evaluated Aii 410 (circle). (B) 3D projection of rendered morphology of Aii 410 from RC1 data with markings illustrating retinal layers occupied by the cell. (C) TEM section (section 650) with Aiis evaluated in RPC1 circled and labeled. (D) Section adjacent (section 649) to the TEM section in C labeled for glycine, medium levels of glycine are consistent with Aii classification. (E) Overlay of images in C and D. (F) 3D projections of rendering of morphology of the 5 Aiis characterized in this study with layers of retina annotated.

In addition to the similar glycine signature of Aiis in RPC1, the gross morphology of the Aii is comparable between the two datasets and consistent with the wider literature of Aii morphology. Namely, the Aiis in RPC1 retain clearly defined lobular and arboreal dendrites characteristic of the Aii cell type [14, 39–41] (Figure 1). The Aii’s soma is located in the inner nuclear layer, directly adjacent to and partially pushing into the inner plexiform layer (IPL), wherein its dendrites stratify in both the OFF and ON layers. The lobular dendrites of the Aii extend within the OFF layer, with a distinct pattern of narrow branches with interspersed, large, vesicle-rich lobules. The arboreal dendrites of the Aii project from the waist of the Aii deep into the ON layer of the IPL, stratifying in the lowest plexiform layers neighboring the ganglion cell layer.

### 1.2 Aii Core Motifs

As described in the introduction, the Aii connectivity has been extensively characterized in the healthy retina [13, 16, 36, 42, 43]. Based on this, we chose to evaluate primary motifs, conserved across mammals, that are characteristic of Aii connectivity within the vertical pathway (Figure 2). Accordingly, we evaluated the ribbon inputs onto the Aii from the broad classes of OFF-cone bipolar cells (CBas), ON-cone bipolar cells (CBbs), and rod bipolar cells (RodBCs). Additionally, we evaluated the gap junction contributions made between each of these cell types and Aiis, as well as between coupled Aiis.

**Fig 2.**
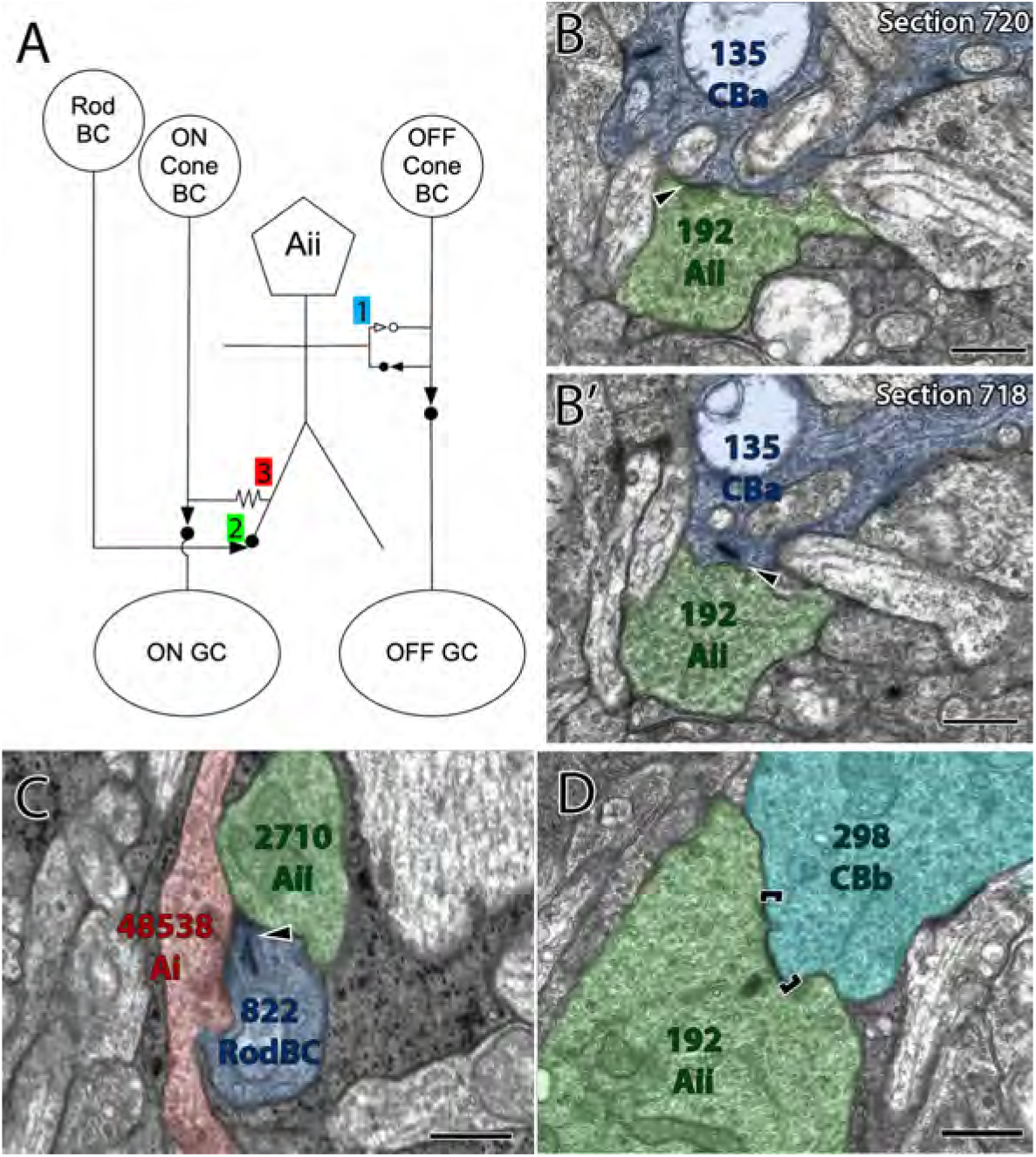
Canonical Aii Network Motifs are preserved in RPC1. (A) Network diagram of simplified Aii connectivity for passage of information of the primary rod pathway. 1: reciprocal synapses made with OFF-cone bipolar cells (CBa), 2: ribbon inputs from rod bipolar cells (RodBCs), 3: gap junctions with ON-cone bipolar cells (CBb). (B) Representative reciprocal synapse illustrating glycinergic conventional synapse from an Aii cell onto a Cba, and (B’) a ribbon synapse from that same CBa onto the same Aii a few sections away. (C) Black arrow highlights ribbon synapse from a RodBC onto a Aii (D) Gap junction between an Aii and a CBb (represented by brackets). *Scale bars: 500nm*

### 1.3 Motif Frequency

Our first metric evaluating the impact of retinal degeneration on Aii motifs was to analyze the synaptic frequency of ribbon input to the Aii and the number of gap junctions made by the Aii with its neuronal partners. Each synaptic pairing described is classified by motif and summarized below. All percentage frequencies are calculated by their percentage of all synapses of that type aggregated by volume and accompanied by the mean and standard deviation of the individual cells involved in the motif. The purpose of presenting both values is to describe the impact of rewiring on the inner retina both globally, by evaluating the motif by volume, while the per cell mean and standard deviation demonstrates the level of variability.

#### 1.3.1 Ribbon Inputs

The first component of input frequency we evaluated was the ribbon inputs by the various classes of bipolar cells (Table 1). Although total ribbon input to Aiis is decreased in in RPC1 (RC1 total ribbon inputs per Aii: 116±12.71, RPC1 total ribbon inputs per Aii: 73.2±11.12), proper lamination is preserved and all bipolar cell superclasses (CBa, CBb, and RodBC) are found within the dataset (Figure 3A-D).

**Table 1.** Ribbon Input by Cell Type.

| Volume | Aii Target | BC |  | CBa |  | CBb |  | RodBC |  |
| --- | --- | --- | --- | --- | --- | --- | --- | --- | --- |
|  |  | % of Total Ribbon Contributions | Ribbon Count | % of Total Ribbon Contributions | Ribbon Count | % of Total Ribbon Contributions | Ribbon Count | % of Total Ribbon Contributions | Ribbon Count |
| RC1 | 410 | 0.98% | 1.0 | 9.80% | 10.0 | 2.31% | 3.0 | 89.22% | 91.0 |
|  | 514 |  |  | 13.85% | 18.0 | 3.77% | 4.0 | 83.85% | 109.0 |
|  | 2610 |  |  | 18.87% | 20.0 | 0.79% | 1.0 | 77.36% | 82.0 |
|  | 3679 |  |  | 15.87% | 20.0 |  |  | 83.33% | 105.0 |
| RPC1 | 192 | 1.79% | 1.0 | 21.43% | 12.0 | 17.86% | 10.0 | 58.93% | 33.0 |
|  | 262 |  |  | 21.62% | 16.0 | 14.86% | 11.0 | 63.51% | 47.0 |
|  | 265 | 1.23% | 1.0 | 28.40% | 23.0 | 1.23% | 1.0 | 69.14% | 56.0 |
|  | 1685 |  |  | 30.68% | 27.0 | 5.68% | 5.0 | 63.64% | 56.0 |
|  | 2710 | 2.82% | 2.0 | 11.27% | 8.0 | 2.82% | 2.0 | 83.10% | 59.0 |

**Fig 3.**
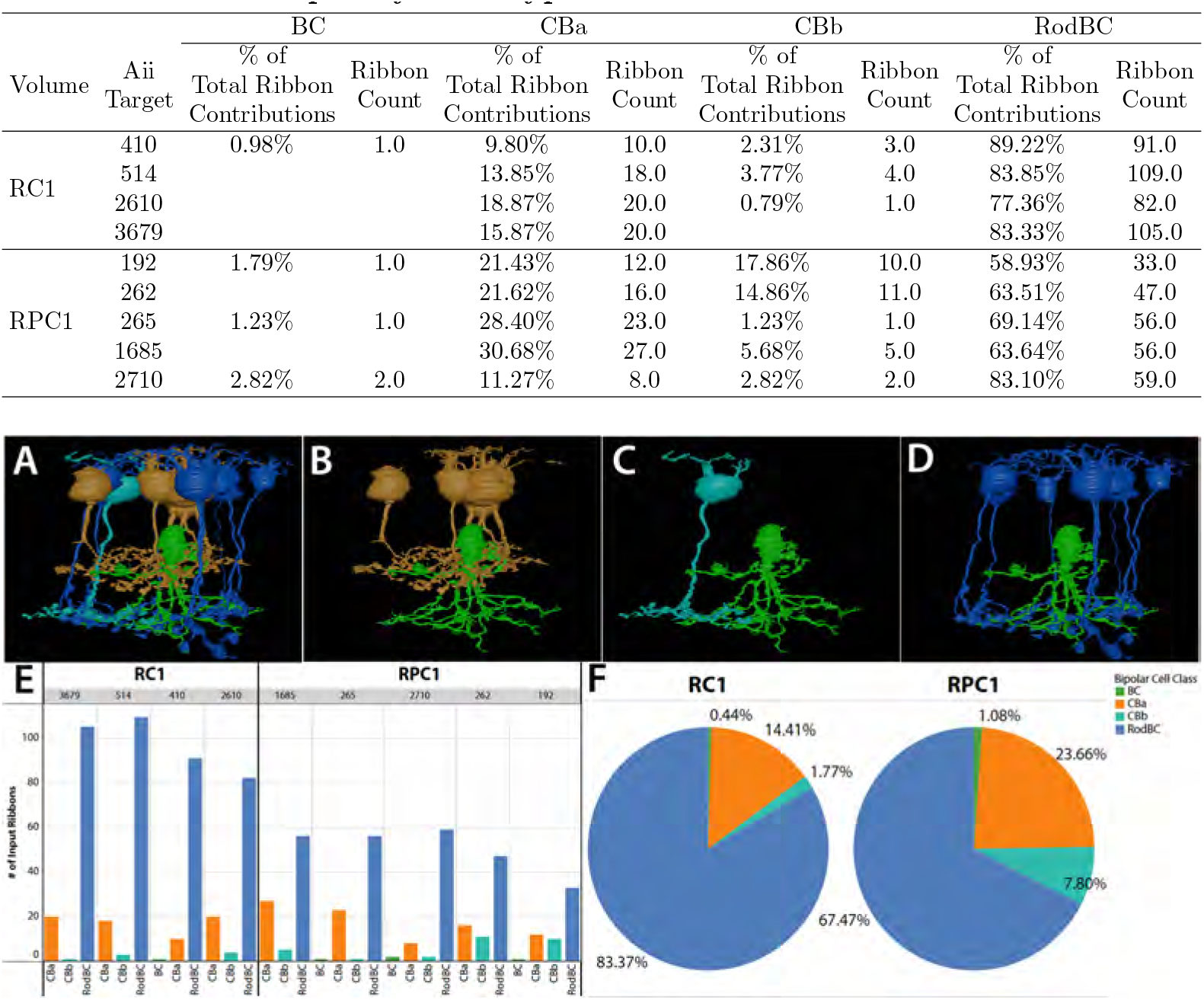
(A) 3D projection images of Aii 265 (green) and the bipolar cells that make ribbons onto it. (B) CBas (gold) presynaptic to Aii 265. (C) CBb (teal) presynaptic to Aii 265. (D) RodBCs (dark blue) presynaptic to Aii 265. (E) Number of input ribbons from each bipolar cell superclass onto each evaluated Aii. (F) Percentage of total ribbon input onto the evaluated Aiis organized by volume.

#### 1.3.2 RodBC →Aii

Extended analysis of previously identified Aiis in the healthy rabbit connectome (RC1) [13]demonstrates a broad number of input RodBCs per individual Aii (10-15). In contrast, the number of RodBCs making ribbons onto Aiis in RPC1 is dramatically decreased to 6-10 RodBCs per Aii. When tabulating numerical ribbon input from RodBCs onto Aiis, we find the total number of ribbons from RodBCs varies between 81 to 109 ribbon synapses per Aii cell, making up 83.37% (mean ± std: 83.39% ± 4.65, coefficient of variation (CV) = 0.05) of total ribbon input to Aiis in RC1 (Figure 3E-F). Further amplifying the observed decrease in RodBC ribbon input in RPC1, the total number of ribbons from RodBCs onto Aiis is reduced to 33-59 ribbons per Aii. This decrease, however, is not uniform across Aiis with the percentage of RodBC ribbon input onto Aiis in dropping to 67.47%, but coinciding with an increase in coefficient of variation (67.38% ± 9.52, CV= 0.13) (Figure 3E-F).

#### 1.3.3 CBa →Aii

The next most common bipolar cell ribbon input partner to Aiis in the healthy retina are the CBa bipolar cells, which are responsible for transmitting OFF network information to the Aii [44]. In RC1, 6-13 individual CBas are presynaptic to each Aii. These CBas make a total of 10-19 ribbons onto each Aii, constituting 14.41% (14.37% ± 3.38, CV= 0.26) of the Aiis’ total ribbon input (Figure 3E-F). In RPC1, the input from CBas is more variable with a total of 8-29 ribbons coming from 3-11 individual CBas presynaptic to each Aii. Overall, the relative input from the CBa superclass onto the Aiis of RPC1 increases to 23.66% (22.99% ± 8, CV=0.33) of the total ribbon contribution (Figure 3E-F).

#### 1.3.4 CBb→Aii

The last contributor of ribbon synapses onto the Aii comes from the CBb superclass. The majority of synapses between Aiis and CBbs are via gap junctions [36]. However, there are a small number of ribbons from CBbs to Aiis found in the healthy retina [13, 36]. In RC1, CBb ribbon inputs exist at a low rate of 0-4 ribbons per Aii, comprising 1.77% (2.29% ± 1.49, CV=0.65) of total ribbon synapses from up to 3 CBbs (Figure 3E-F). In RPC1, Aiis receive ribbons from as many as 7 CBb partner cells with a wide range of 1-11 ribbons constituting 7.80% (8.47% ± 7.45, CV=0.87) of all ribbon inputs to each Aii (Figure 3E-F).

#### 1.3.5 Aii Gap Junctions

Aberrant gap junctions between RodBCs and Aiis were first ultrastructurally described in the RPC1 volume [35]. No gap junctions have ever been directly observed or described in the literature between RodBCs and Aiis in the adult mammalian retina, leading us to conclude their emergence in the RPC1 volume is secondary to retinal plasticity events and retinal remodeling induced by rod degeneration. Here, we expand upon previous efforts to evaluate the gap junction motifs in the healthy and degenerate retina and explore the relative contributions by each gap junction partnership (Figure 4).

**Fig 4.**
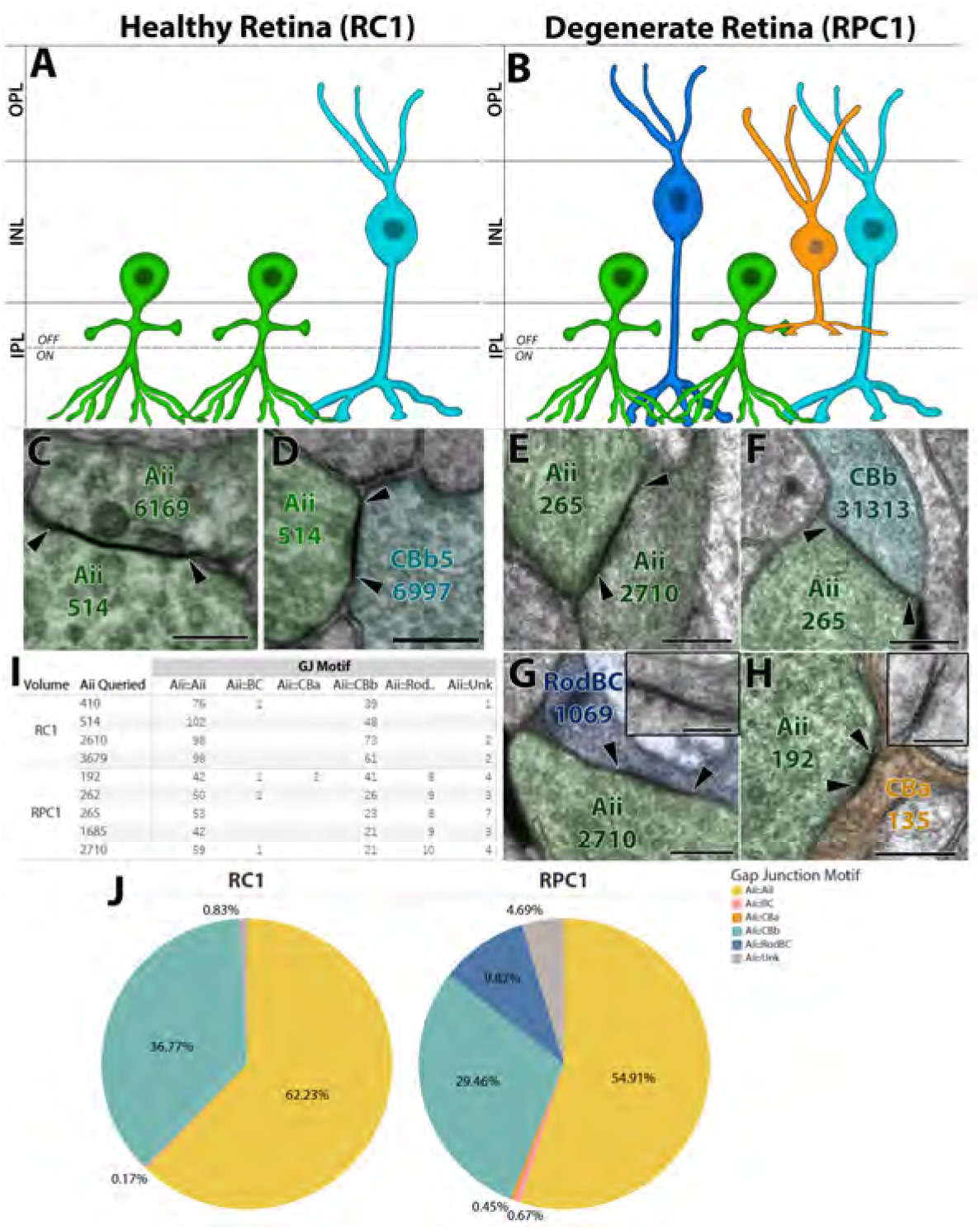
(A) Diagram illustrating cell classes coupled with Aiis in the Healthy retina. (B) Diagram illustrating cells coupled with Aiis in the degenerate retina. (C-D) Gap junctions of both motifs present in the healthy retina, (C) Aii :: Aii and (D) Aii :: CBb. (E-H) Representative gap junctions present in RPC1 (E) Aii:: Aii, (F) Aii :: CBb, (G) Aii :: RodBC, *inset image:* Higher magnification recapture @ 25k with 20 degree tilt, (H) Aii :: CBa *inset image*: High magnification recapture @ 25k with 4 degree tilt. (I) Raw counts of the gap junction motifs per analyzed cell in each volume. (H) Percentage of total number of gap junctions contributed by each motif. (scale bars *Primary panels: 250nm, inset images 100nm)*

##### Aii::Aii

The most prevalent gap junction pairing of Aiis in the healthy retina is with other Aiis [13, 16, 36] (Figure 4C). The raw number of Aii homocellular gap junctions (gap junctions with cells of the same class) observed in RC1 is between 76 and 102 Aii::Aii gap junctions (Figure 4I). The contribution of the Aii::Aii motif to the total number of gap junctions is 62.2% (62.62% ± 4.93, CV= 0.07) (Figure 4J). In RPC1, this motif is reduced to 54.91% of the total gap junctions (Figure 4E), although the cell-to-cell variability is increased (55.19% ± 7.28, CV= 0.13). This is further demonstrated by each individual Aii making 42-59 homocellular gap junctions (Figure 4I).

##### Aii::CBb

The gap junctional coupling between Aiis and CBbs is critical for the connection of rod inputs to the RGC because RodBCs are not presynaptic to RGCs [14, 16](Figure 2A). In RC1, the Aii::CBb pairing is the other prominent gap junction motif of the Aii [13, 36] (Figure 4D). Although a wide raw number of gap junctions of this motif exist per cell (39-73) (Figure 4I), the contribution of this motif is 36.77% of the Aii gap junctions in RC1 (Figure 4J) (36.35% ± 4.64, CV=0.13). In the degenerate RPC1 retina, Aii::CBb gap junctions (Figure 4F) constitute a smaller fraction of the total gap junction population at 29.46% (Figure 4J) with higher variability (29.21% ± 7.55, CV=0.26) and a raw number of 21-41 gap junctions (Figure 4I).

##### RodBC :: Aii

These findings are among the most remarkable in early degenerate retina, providing a pathway for corruption of visual processing. Although the emergence of gap junctions between Aiis and RodBCs are novel to the early stage degenerating retina [35] (Figure 4G), their frequency is remarkably consistent at 8-10 gap junctions per Aii (Figure 74I).

With the addition of these gap junctions to the degenerate network, RodBCs make up 9.82% of all gap junctions made by Aiis in RPC1 (Figure 4J), again with lower variability than many other observed motifs (9.89% ± 1.45, CV=0.15). All RodBCs making gap junctions with Aiis also continue making ribbon synapses onto the same Aiis (Figure 4G).

##### Other Aii pairings

We also find within RPC1 the rare occurrence of Aii::CBa gap junctional coupling, as evidence by 2 instances of this pairing (Figure 4H). In these instances, branches of a single CBa (Cell# 135) were found to make small gap junctions with the waist region of a neighboring Aii (Cell# 192). These are another motif that has not been observed within the healthy retina, however it was exceptionally rare constituting only 0.45% of all gap junctions (Figure 4J), in contrast with the aberrant gap junctions observed with RodBCs.

While the gap junction motif percentages are certainly altered by the addition of the RodBC::Aii motif in RPC1 (Figure 4J), part of the changes in relative contributions of the Aii::CBb and Aii::Aii motifs is the increase of unknown partners (Figure 4I). Unknown partners arise due to branch fragments that cannot be followed, generally due to rapidly going off the edge of the volume. In RC1, unknown partners contributed 0.83% of all gap junctions with Aiis. In RPC1, this number increased to 4.69% (Figure 4J), likely due to the smaller diameter of the RPC1 volume. Additionally, there were 3 instances of bipolar cells that could not be identified. These incomplete classifications, however, only make up 5.36% of all of the gap junctions made by Aiis in RPC1.

### 1.4 Aii Synaptic Weighting

Extending our understanding of network inputs based on synaptic tabulation from specific cell classes, we explored how potential differences in synapse size affect synaptic weighting, or the relative strength of contributions, of each motif in the healthy and degenerate retina. To better estimate the potential synaptic weight, we directly measured the area of gap junctions and post-synaptic densities (PSDs) opposing presynaptic ribbons. The concept that gap junction plaque size is a contributing measure of strength is previously established [36]. However, it is not possible to evaluate pore conformation, phosphorylation state, nor rectification from TEM analysis alone, therefore this gap junction synaptic weight should be regarded as the maximum potential synaptic weight. Cortical brain spine size is strongly correlated with synaptic strength [45], while the size of the readily releasable pool is not [46], and spine size has been previously used by others as a measure of relative input [47]. In the retina, spines are not formed at PSD sites necessitating another metric of strength. Given the spine volume being proportional to the area of the PSD [48], we chose to measure the area of the PSD opposing ribbon-type synapses as a metric of synaptic weight (Equation 1), where *n* = number of sections PSD density extends across, *l* = length of density on given section, *t* = thickness of section (set at 70nm based on section thickness of volume).

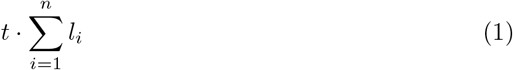

#### 1.4.1 Ribbon Synapse Strength

As described above, we evaluated the area of all PSDs on Aiis opposing a ribbon density, allowing us to approximate the strength of bipolar cell glutamatergic input from each class (Figure 5). This was done as an extension of the ribbon count metrics used in the previous section to determine whether the observed alterations in ribbon input counts by bipolar cell class to the Aii is representative of true alterations in strength of glutamatergic drive.

**Fig 5.**
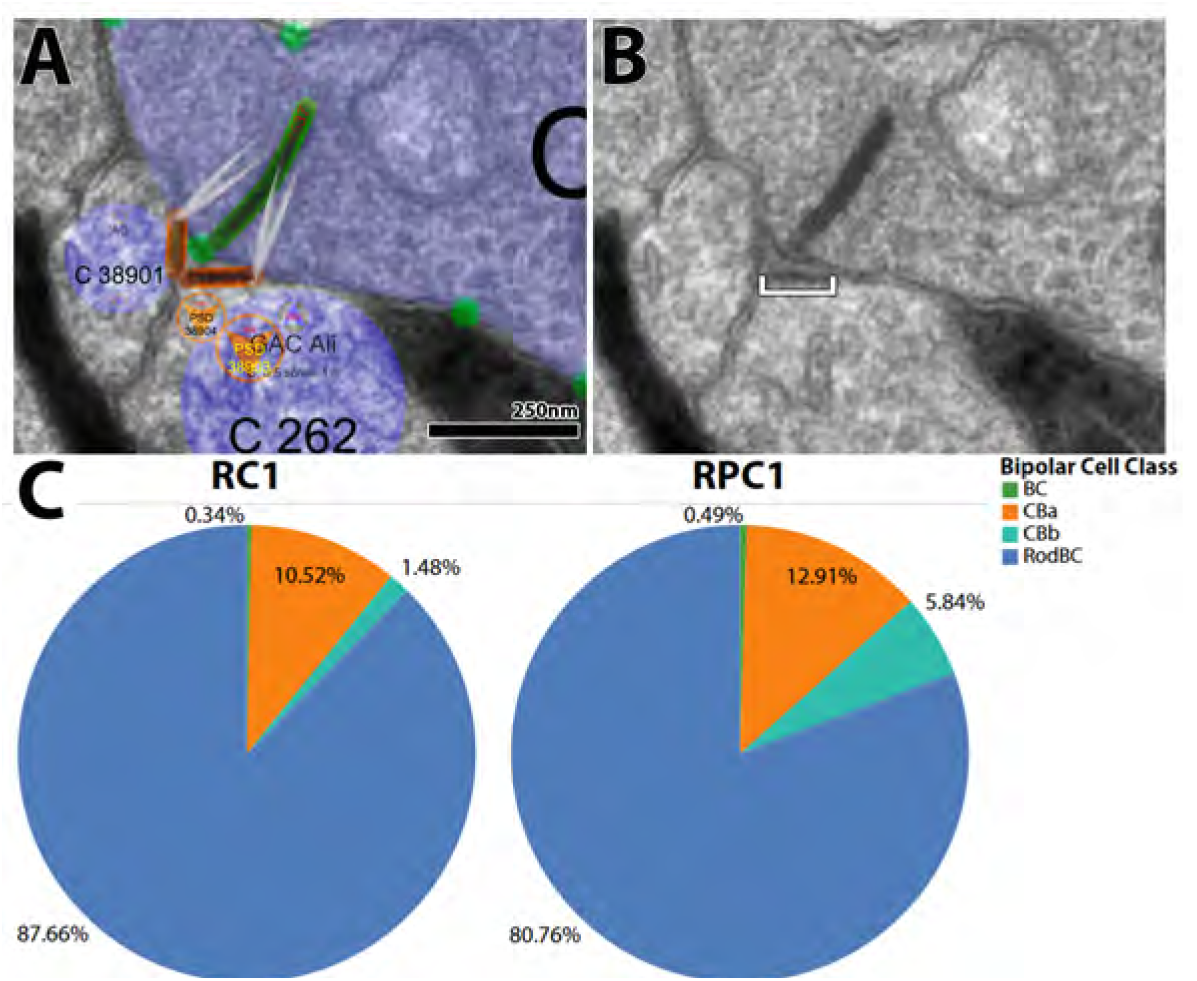
(A) Annotated section demonstrating use of annotations to determine size. Green bar: ribbon, orange bar: annotated PSD, orange circles: references to annotations on adjacent sections. (B) Same section as shown in panel A without annotations. PSD is indicated by white bracket. (C) Percentage of contribution by each motif per volume as calculated by total gap junctional area.

#### 1.4.2 RodBC →Aii

Consistent with the raw counts of RodBC ribbon synapses onto Aiis of the healthy retina, quantification of PSD area indicates that RodBCs are the greatest contributor to Aiis of all BCs in both the healthy and degenerate retina (Figure 5C). We find that RodBC-opposed PSDs account for 87.66% of all ribbon PSD area in RC1, with the average PSD being 0.03µm^2^ ± 0.017. This input demonstrates the total RodBC input strength is roughly 4% higher than indicated from raw count alone. In RPC1, RodBC-opposed PSDs are substantially larger than in RC1 at 0.04µm^2^ ± 0.025 accounting for 80.76% of all ribbon PSD area, a full 13% greater input strength than is indicated by ribbon synapse count alone. The PSD sizes in RPC1 potentially compensate for the lower numbers of ribbon inputs, which may help to equalize the relative strength of RodBC input to the Aiis. We model this effect below.

#### 1.4.3 CBa →Aii

Following the overwhelming input from RodBCs, the next greatest ribbon input comes from the CBa class. In RC1, CBa inputs make up 10.52% of the total ribbon PSD area with an average PSD size of 0.02µm^2^ ± 0.013. In RPC1, CBa-opposed PSDs make up 12.91% of the total ribbon PSD area with an average area of 0.018µm ± 0.014. In both cases, the small size of the PSDs demonstrates a decrease in network weight from that indicated by ribbon-synapse count alone (14.41% in RC1 and 23.66% in RPC1).

#### 1.4.4 CBb→Aii

Lastly, CBb ribbon inputs onto Aiis are by far the smallest contributor of glutamatergic input. In RC1, the CBb-opposed PSDs make up 1.48% of all ribbon PSD area with an average size of 0.024µm^2^ ± 0.018. This is comparable to the 1.77% input calculated from ribbon inputs alone. In RPC1, the CBb-opposed PSDs make up 5.84% of the total ribbon PSD area, down from 7.8% calculated from ribbon input count alone, with an average area of 0.018µm^2^ ± 0.014. The contributions in RPC1, however, are highly variable between individual Aiis. CBb-opposed PSDs may make up as little as 0.36% of all ribbon input (seen on Aii Cell # 265), or as much as 16.46% (seen on Aii Cell # 262).

### 1.5 Aii Gap Junction Size

As described by Sigulinsky et al. 2020 and Marc et al. 2014 [13, 36], Aii gap junctions are exceedingly precise, both in coupling partners (as evidence by the lack of gap junction formation despite opportunity in specific bipolar cell classes) and in gap junction plaque size [36]. Gap junction size has been shown to correlate with strength [49, 50], and is tightly associated with retinal gap junction partner motifs [36]. Here, we build upon the coupling rules established in RC1 [13, 36], and report the relative contributions of gap junctions in the Aii network of RPC1 by cell superclass partner (Figure 6).

**Fig 6.**
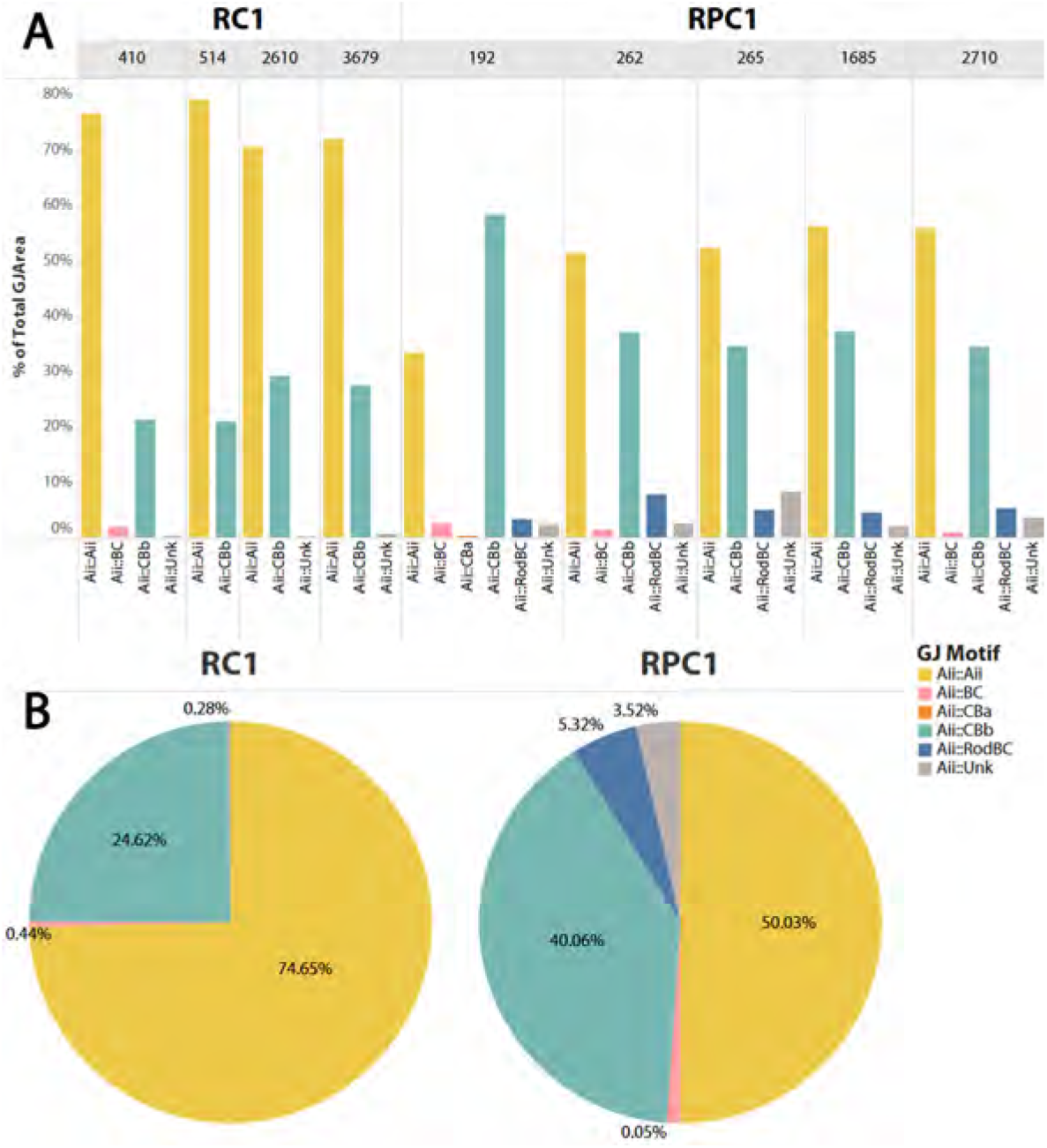
(A) Percentage of gap junction area contributed by each motif separated by cell and volume. (B) Percentage of total gap junction area contributed by each motif per volume.

#### 1.5.1 Aii::Aii

Consistent with gap junction motif frequency determined by raw count and previously published reports [13, 14, 16, 51], the strongest coupling motif remains the Aii in-class coupling with other Aiis (Figure 6). In RC1, the area of gap junctions between Aiis make up 74.65% of all gap junctional area with an average area of 0.12µm^2^ ± 0.1. In RPC1, this motif is still the strongest contributor to the Aii coupling network, but has dropped substantially to 50.03% of all gap junctions by area. This is further highlighted by the average area being substantially smaller at 0.05µm^2^ ± 0.04.

#### 1.5.2 Aii::CBb

As described in the previous section, the Aii::CBb motif is the other primary gap junction motif found in the Aii gap junction network. In RC1, this motif accounts for 24.62% of total gap junctions by area, with an average area of 0.065µm^2^ ± 0.06. In RPC1, however, the contribution of the Aii::CBb motif is dramatically increased to 40.06% of total gap junctional area and an average area of 0.073µm^2^ ± 0.054.

#### 1.5.3 Aii::RodBC

The emergence of the RodBC::Aii gap junction motif in RPC1 was the major impetus for this analysis. As previously noted, there are no gap junctions of this motif in RC1, nor have any ever been found in a healthy adult mammalian retina, and represents and entirely new architecture, albeit pathological, but predictable and stereotyped. In RPC1, this motif accounts for 3.29-7.9% of the gap junctional area made by individual Aiis, with 5.32% contribution to total gap junction area. The average Aii::RodBC gap junction in RPC1 is 0.029µm^2^ ± 0.02.

#### 1.5.4 Other Aii pairings

Lastly, the emergent gap junctions between a CBa (Cell # 135) and a Aii (Cell # 192) are exceedingly small. Although this motif has not been observed in the healthy retina, in RPC1 these make up only 0.05% of the total gap junctional area.

The unknown and incompletely identified gap junctions of RC1 contribute a combined 0.72% of the total gap junction area in RC1. In RPC1, these incomplete partnerships account for 4.57% of the total gap junction area of the analyzed cells. The unidentified partners are likely higher in RPC1 due to the smaller diameter of the volume.

## 2 Simulation Results

Combined, the above description allowed us to investigate the specific network changes associated with rewiring in early RP, specifically the rewiring of the neural circuitry contributing to degeneration-unique signaling patterns. Here, we evaluate the impact of these altered networks on the retina’s electrical activity. The network model is constructed using realistic, multi-compartmental morphologies and cell-specific membrane properties (for further detail, see supplemental methods S1).

### 2.1 Baseline

First, the baseline light response of each retinal cell type was simulated to establish a ground truth by excluding the aberrant synaptic connections found in the RPC1 volume. The differences between baseline and degenerate networks are summarized in Table 4. Responses of dark-adapted RodBCs, CBbs, Aiis, and the RGC were evaluated during light stimulation (Figure 7). ON bipolar cells exhibited sustained depolarization due to graded ribbon synaptic input from photoreceptors. Consequently, the RGC generated action potentials during the depolarization phase of CBbs. Aiis displayed both transient and sustained EPSP components. The initial transient spike was driven by rod input and disappeared when the rod photocurrent was set to 0 pA. This behavior is consistent with experimental recordings of RodBC–Aii ribbon synapses, where sustained inward calcium currents evoke a transient EPSP spike before reaching steady-state depolarization [52]. The sustained depolarization of Aiis persisted as long as CBbs remained depolarized, in agreement with patch-clamp recordings obtained using the same light flash protocol. The RGC model reproduced impulse response characteristics reported previously [53, 54]. At full cone saturation (20 pA), the simulated peak firing rate reached 120 Hz, within the range reported in patch-clamp studies of light-evoked RGC responses [55, 56]. We defined RGC firing frequency as the visual output and compared it between baseline and degenerate networks, hypothesizing that changes in RGC firing rate reflect functional impairment during early-stage RP.

**Fig 7.**
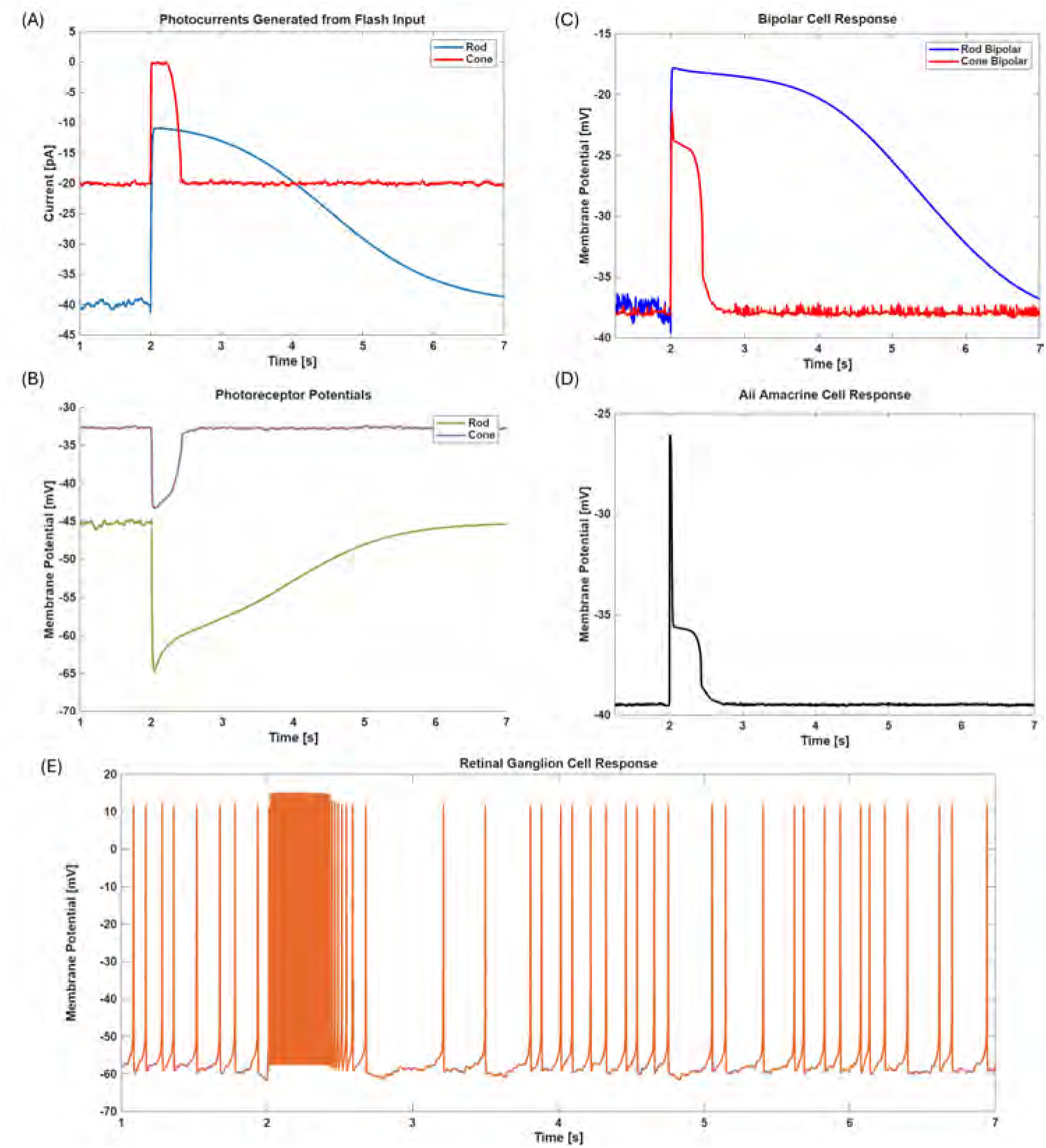
Changes in the EPSP of retinal neurons following light stimulus. Cells begin at rest (dark-adapted) and the stimulus is applied at the 2-second mark. (A) The photocurrent model for rod and cone photoreceptors due to a light flash protocol. The photocurrents sit at a negative level at rest, accounting for the dark current, and have random noise to model the spontaneous activity of the RGCs. (B) The photoreceptor inner segments become hyperpolarized after stimulus, limiting the release of glutamate neurotransmitters. (C) The bipolar cells become depolarized as the rate of glutamate binding to mGluR6 is reduced. The dark noise is reflected as minor oscillations of the EPSP, not exceeding 2 mV in amplitude. (D) The Aii EPSP contains two components: a transient fast-acting spike due to excitatory ribbon synapses with the RodBC, and sustained depolarization due to gap junctions with the CBb. (E) The RGC demonstrates spontaneous, non-rhythmic firing at rest and high-frequency, rhythmic firing as long as the cone photocurrent persists.

### 2.2 Early-Stage Degeneration

The aberrant gap junctions between RodBCs and Aiis are based on the documented connections characterized above in RPC1, where 27 were identified across the 14 RodBCs and 5 Aiis. These gap junctions are exclusive to RPC1 and have thus far only been described during early-stage degeneration. By adding these aberrant gap junctions into the baseline network model, the impact of early-stage degeneration on each cell’s EPSP is compared with those of the baseline.

#### 2.2.1 Dark-Adaptation

The RGCs exhibit spontaneous activity during rest, which becomes oscillatory and hyperactive during early-stage degeneration. The dark-adapted resting state of the RGC was was simulated across four seconds and the firing rate was compared between baseline and degenerate networks (Figure 8A). The average firing rate between the baseline and degenerate RGCs were 6 Hz and 10 Hz across 4 seconds, respectively, amounting to ~4 Hz difference during spontaneous resting activity. These values are consistent with experimental recordings of degenerate retinas in the rd1 and rd10 mice [57].

**Fig 8.**
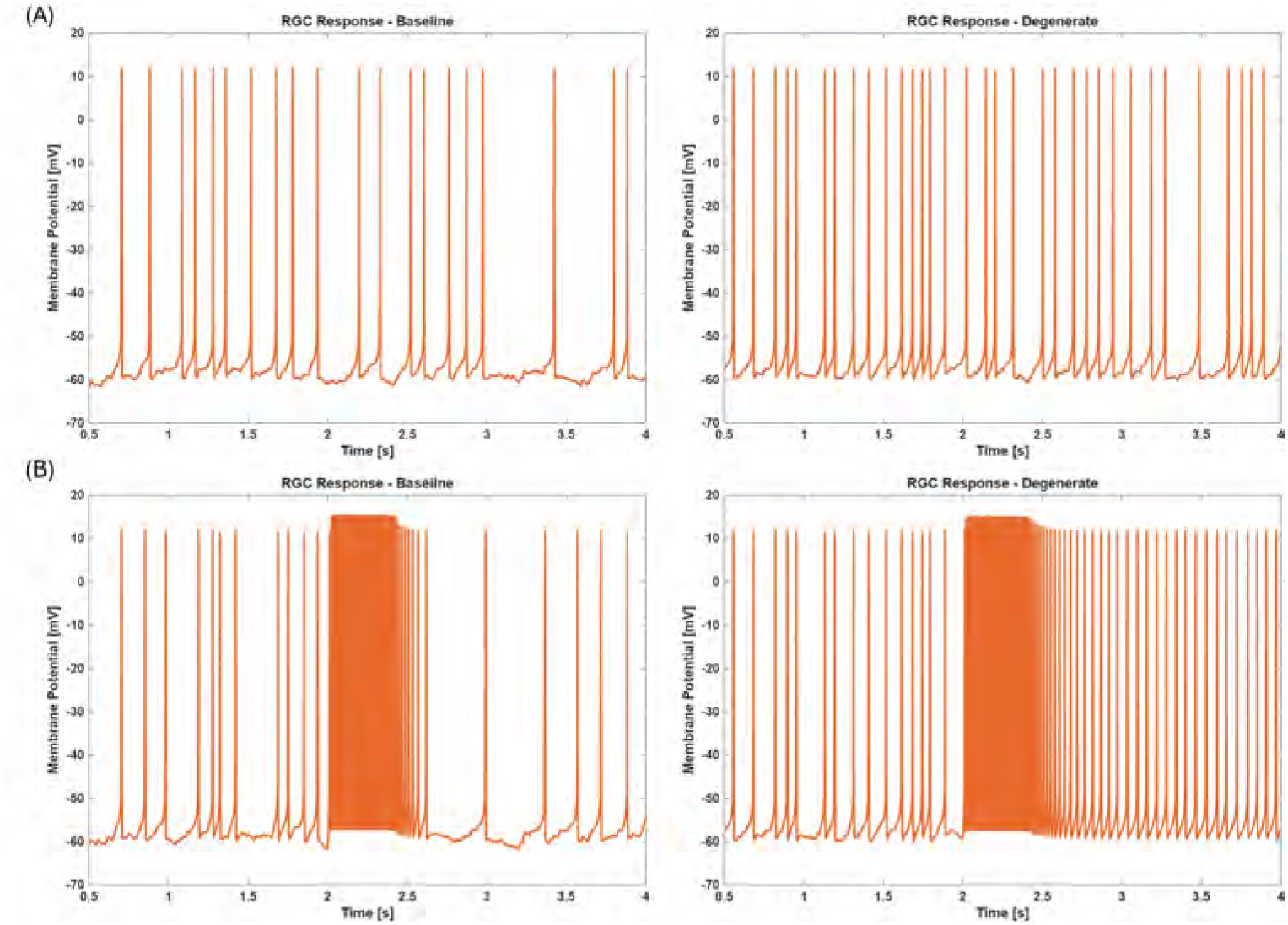
The gap junction rewiring between RodBC and Aiis impact the firing profile of the RGC. (A) The spontaneous firing of the degenerate RGC is faster and more rhythmic in the abscence of light stimulus. Baseline firing rate is 6 Hz on average whereas degenerate rate is 10 Hz across 4 seconds of simulation (B) The firing profile shows significant difference after the stimulus (3 - 4 s). The baseline rate is the same as in (A), whereas the degenerate rate becomes 16 Hz and rhythmic. The firing rate of either RGC during stimulation is 120 Hz.

#### 2.2.2 Cone-Saturating Light Response

Patch clamp recordings are typically conducted on dark-adapted retinas, where brief light pulses are used for stimulation. A light pulse was applied at t = 2 seconds to simulate the natural light response, as described in section 2.6.3. Key trends emerged across all retinal neurons in the degenerate network, where the resting membrane potentials became higher than those of the baseline network (Figure 9). Before light stimulation, the resting membrane potential of the degenerate Aii (*V*_*m*_) is 1 mV higher than that of the baseline Aii. Notably, this difference becomes 3.2 mV post-stimulation. In the healthy retina, the RodBC-Aii synaptic connectivity is mediated solely by AMPA-type glutamate receptors [58], generating a transient spike on the Aii EPSP (Figure 7D). However, the presence of gap junctions creates an electrical coupling that further depolarizes the Aiis while the rod photocurrent persists. As Aiis and CBbs are also electrically coupled, the change in *V*_*m*_ is reflected on the CBb in the same ratio (Figure 9). The CBbs depolarize up to 1.8 mV from their baseline resting EPSP as a result. The RGC fires at a greater rate in the degenerate retina because of this difference, especially during the post-stimulation period (Figure 8B). It is evident in our model that the depolarization of the retina is the driving mechanism of hyperactivity associated with degeneration. This is a consequence of the electrical coupling that does not occur in the baseline network, leading to a mixing of rod signals into the cone pathway for an extended period.

**Fig 9.**
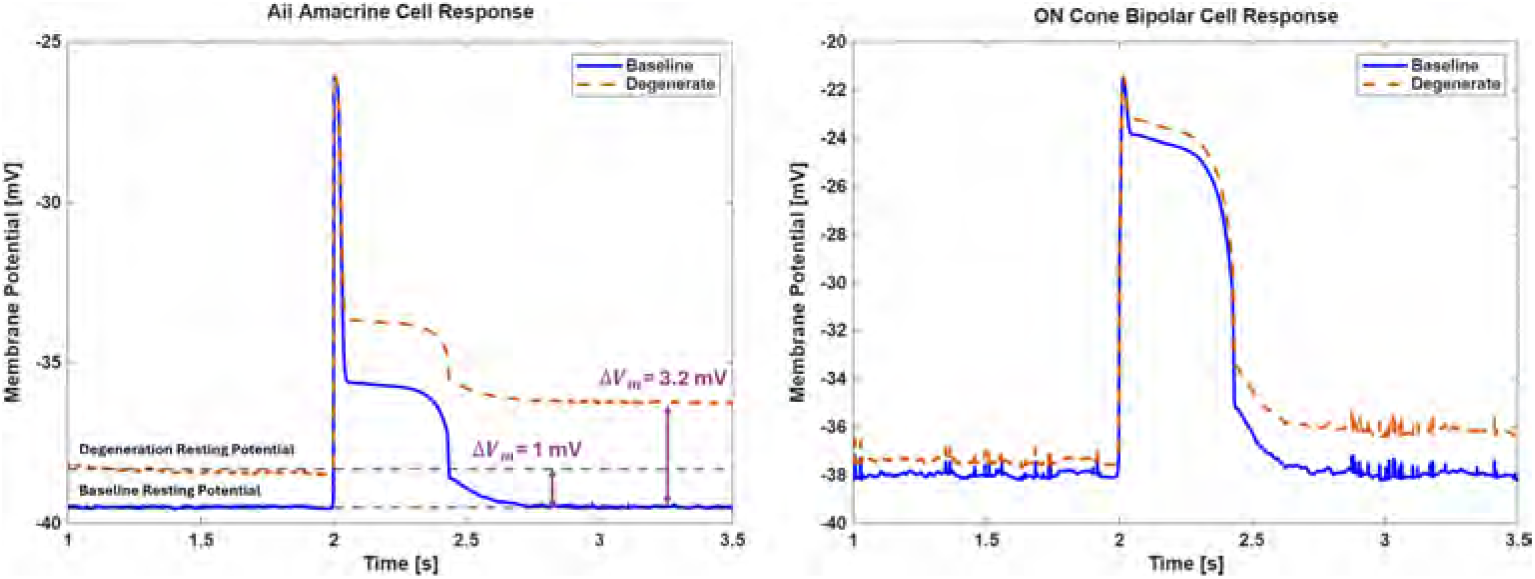
Changes in membrane potentials of the amacrine and bipolar cell types following a saturating flash of light input shown in Figure 7A at the 2-second mark. A depolarization effect is observed on the resting membrane potentials in the degenerate network. The depolarization is stronger after the 0.5 s stimulation pulse, up to 3.2 mV higher than the baseline for Aiis and up to 1.8 mV for CBbs.

### 2.3 Model Parameter Sweep and Impact on the Output

#### 2.3.1 Number of Rod Inputs per RodBC

In section S1, we highlight varying reports regarding the number of rod inputs received by each RodBC in the healthy rabbit retina. Whereas this count was established at 30, based on referenced modeling studies, additional evidence suggests the possibility of it reaching as high as 100 [59]. We note that the ONL thickness had reduced by 50 percent in the RPC1 and the degenerating rods that are still alive may have already lost their synapses with the RodBCs. Consequently, accurately estimating the intact connections within the imaged volume became challenging. To address this uncertainty, we performed simulations, sweeping through 0 to 90 rod inputs per RodBC in the degenerate retina model, using intervals of 15 rods/RodBC (Figure 10). The RGC output was evaluated in two regimes: the peri-stimulation between t = 2 s and t = 2.5 s, and the post-stimulation between t = 3 s and t = 4 s. The peri-stimulation RGC firing rate did not vary with increasing rod inputs per RodBC, remaining at around 120 Hz, as the cone photocurrent dominates its output. In contrast, we observed a logarithmic pattern in the RGC firing rate in the post-stimulation regime while the cone photocurrent was zero, indicative of the effect of an increasing number of rod inputs. The increase in the rhythmic firing is thus attributed to the sustained feeding of rod signals into the cone pathway, resulting in depolarization of the CBbs post-stimulus.

**Fig 10.**
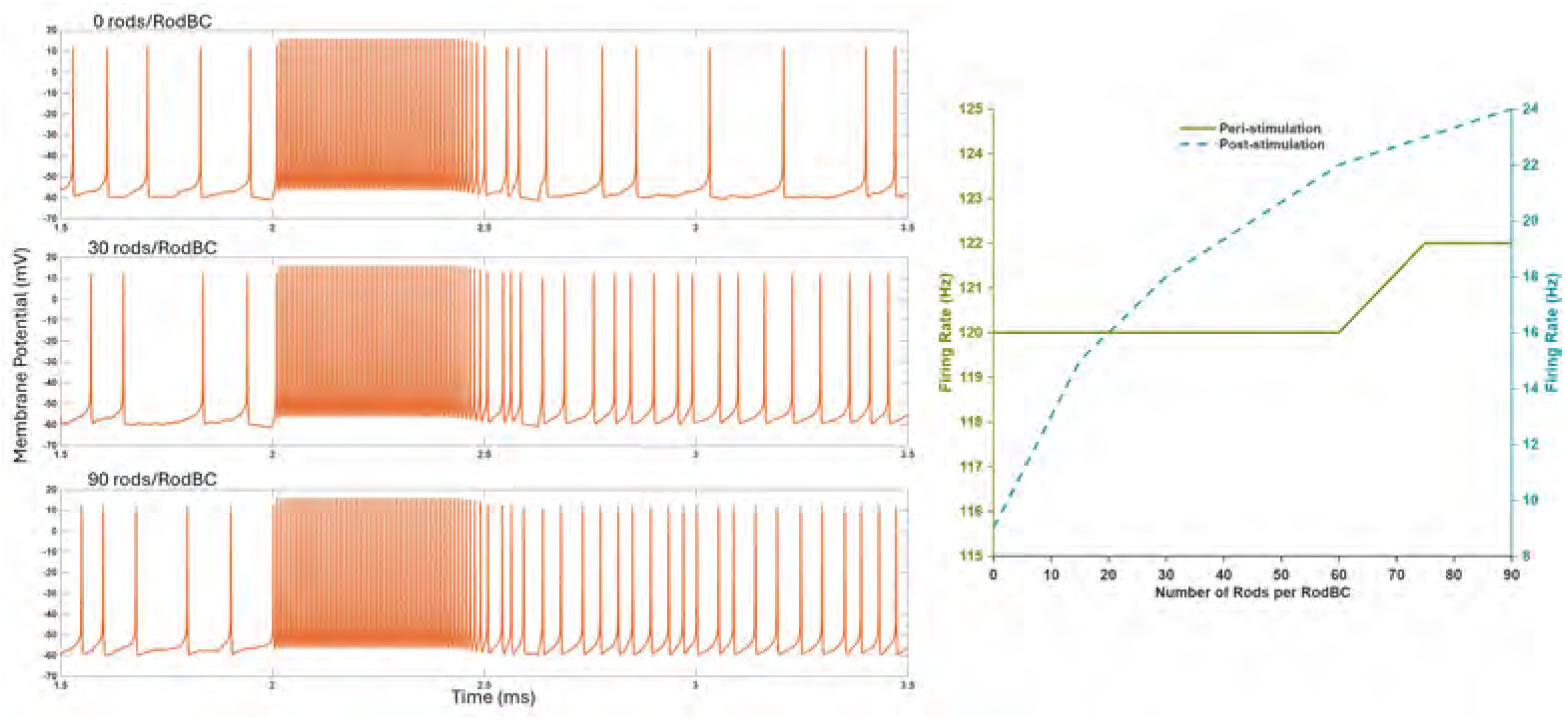
Left: The RGC membrane potential in the degenerate retina plotted when no rods, 30 rods and 90 rods were connected to each RodBC. A sustained rhythmic firing was observed after the end of cone photocurrent for 30 and 90 rods per RodBC whereas non-ryhthmic spontaneous firing was observed when no rods were connected. Right: The peri-stimulation (saturated cone photocurrent) RGC firing rate remained at 120 Hz whereas the post-stimulation (zero cone photocurrent) RGC firing rate between t = 2.5 s and t = 3.5 s followed a logarithmic relationship with respect to the number of rods per RodBC. The rhythmic activity occurred only in the presence of rod signals, approaching saturation around 25 Hz as the rod number increased.

#### 2.3.2 Gap Junction Conductance

Finally, we simulated the contribution of all three network topologies of gap junctions to the RGC output by using a greater conductance value. How the electrical coupling between each cell type affects the RGC output is quantified in Table 2. Increasing the conductance of all gap junctions in the model from 200 pS to 700 pS had minimal effect on the post-stimulation RGC firing rate, at about 1 Hz. The blockage of different gap junction groups by setting their conductance to zero, confirmed the role of RodBC gap junctions in the rhythmic hyperactivity of the RGC. The firing rate remained elevated at 16 Hz when Aii::Aii or Aii::CBb gap junctions were blocked, demonstrating that the RodBC connections drove the hyperactivity.

**Table 2.**
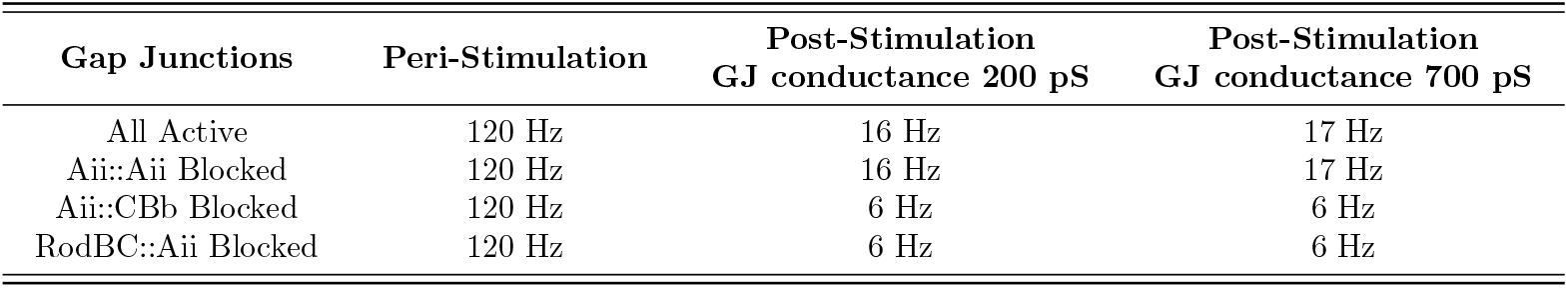
The impact of blocking different gap junction groups and increasing their conductance on the RGC firing rate peri-stimulation (cone photocurrent active, 2*s < t <* 2.5*s*) and post-stimulation (cone photocurrent zero, *t >* 2.5*s*).

## Discussion

Pathoconnectomics is an emerging field that describes aberrant neural connectivity in networks arising during degeneration. In this study, we further investigated the impact of retinal degeneration on the network topologies of Aii cells in the RPC1 pathoconnectome by comparing them with the healthy rabbit connectome RC1 and modeling the effects of these altered topologies. RC1 has been continuously annotated for over 15 years, providing extensive information about the healthy Aii networks [13, 36, 60], and supported by decades of prior work [16, 41, 51, 61, 62]. The primary Aii network components evaluated in this study are the ribbon inputs from the 3 superclasses of bipolar cells (RodBC, CBa, and CBb) and the gap junctional motifs generated between Aiis and their partners, focusing upon the impact of retinal degeneration. Additionally, we expanded our evaluation of the network changes observed in the degenerate retina by including synapse size as a metric of synaptic weight. The graphical representation of this network topology is presented in Figure11. Furthermore, through modeling, we contextualize how Aii network changes potentially underly observed disease mechanisms.

### 1) Rewiring Affects RGCs Response During Low-Light Conditions

In this study, we constructed a retinal network model incorporating the network topologies identified from the RPC1 volume and simulated the impact of aberrant RodBC::Aii gap junctions on the RGC output. The results show that these aberrant gap junctions lead to hyperactivity in RGCs. Simulations further indicate that the observed hyperactivity primarily depends on the rod pathway (Figure 10). Current evidence from pathoconnectomics suggests that RodBCs form pathological gap junctions with Aiis prior to complete photoreceptor loss and that Aiis alter their synaptic contacts early in the degeneration process. If rewiring precedes rod death, it may be possible to detect disease onset through changes in RGC activity before symptoms of diminished night vision appear, as shown in (Figure 8). This feature is particularly relevant for early intervention strategies aimed at preserving photoreceptors [63].

### 2) Network Depolarization via Aberrant RodBC - Aii Gap Junctions Drives Rhythmic RGC Firing

Previous work has shown that aberrant electrical activity of elevated and rhythmic spiking patterns is present in the RGCs of numerous retinal degeneration models [64]. This hyperactivity reduces the fidelity of signal being transmitted from the retina to higher visual processing centers of the brain [65], potentially contributing to altered phosphene perception and loss of visual acuity characteristic of retinal degeneration [66]. The origin of this aberrant spiking has been largely attributed to the Aii network, specifically the Aii-CBb connections [67, 68]. Furthermore, the aberrant RGC spiking patterns are attributed to gap junctions via Cx36 coupling [65] and gap junction phosphorylation states [69]. Prior modeling efforts based on the known coupling motifs of healthy retina (homocellular Aii::Aii gap junctions and heterocellular CBb::Aii gap junctions) also attribute rhythmic RGC firing to intrinsic cellular mechanisms and Aii::CBb coupling [57, 67]. Our results provide a clear explanation of the mechanism of oscillatory potentiation in degeneration. Through pathoconnectomics, we show that RodBC::Aii coupling drives a stronger and persistent coupling of the rod pathway with the cone pathway, thereby increasing the RGC activity through the CBbs. Our model demonstrates that a 4-10 Hz oscillation emerges after a 1-3.2 mV increase in the resting potential of Aiis (Figure 9), aligning with the findings of similar degeneration modeling study where rd1 mouse demonstrated rhythmic activity up to 9 Hz following a 2-3 mV increase in the median Aii [57]. Simulations show that the RodBC::Aii gap junctions induce these oscillations by shifting the basal resting potential of Aiis. Critically, blocking RodBC::Aii gap junctions, while preserving Aii::CBb gap junctions, prevents Aii depolarization at steady state and eliminates the rhythmic RGC firing (Figure 8). Furthermore, depolarization of RodBCs, which produces a transient Aii EPSP in the baseline network (Figure 7D), instead generates a sustained post-transient EPSP in the presence of aberrant RodBC::Aii coupling (Figure 9), as long as rod photocurrent remains active. This elevated potential propagates to CBbs via existing gap junctions, thereby increasing their depolarization level and firing rate of the RGC. Together, these findings support the hypothesis that the elevated steady-state cell membrane potentials during rod activation drive the rhythmic RGC firing in the degenerate retina. Our conclusions are consistent with a similar computational study of later-stage retinal degeneration [70], which reports that depolarization of the ON-pathway and the Aiis produces patterned rhythmic activity, as cone photoreceptor density declines. The authors note that rod pathway contributions may be underestimated in their model and suggest that rod-driven oscillations may occur in earlier stages. Our results support this hypothesis, demonstrating that such oscillations can arise specifically from aberrant RodBC::Aii coupling during early-stage degeneration.

### 3) Aberrant Activity Scales with the Intensity of Rod Signal and Gap Junction Conductance

Our simulations addressed topological uncertainties in the pathoconnectome data. Due to the necessary classification of many photoreceptor synapses as indeterminate, and given our focus on rod pathway degeneration, we performed simulations to evaluate how rhythmic RGC firing depends on rod signal strength. The results indicate that the rod pathway modulates Aii depolarization and serves as the primary driver of RGC hyperactivity (Figure 10). Increasing rod convergence toward the upper bound reported in rabbit retina (approximately 100 rods per RodBC) results in saturation of RGC firing at approximately 25 Hz. The simulated range of rhythmic firing (10–25 Hz) agrees with experimental whole-cell recordings of Aiis and RGCs in degenerated retinas [65, 71–75]. The awareness of the impact of varying rod convergence is valuable in vision research, as coupling and receptive fields of RodBCs and Aiis may exhibit substantial variation based on factors such as light intensity and eye size. For example, a recent study revealed a higher convergence of rods and RodBCs to a single downstream Aii in macaques compared to mice [76]. This increased convergence led to a receptive area twofold greater, maintaining necessary resolution within the same visual angle. Moreover, the same study underscored the impact of light adaptation on the coupling between Aiis, modulating a balance between resolution and sensitivity. Computational models such as presented in this work provide the flexibility to modify parameters between species and produce a more realistic signal processing of the visual output.

Additionally, we investigated the influence of gap junction conductance on neuronal responses, where modulating gap junction conductance had a modest effect on aberrant firing rates, of about 1 Hz (Table 2). However, the implications of variable gap junction coupling between different cell types may become more significant in larger network models with more cells and synapses. Physiologically, a higher gap junction conductance has been suggested to reduce signal noise between adjacent cells, and a coupled network of Aiis can amplify small but correlated rod signals [77, 78]. Simulating a larger network can illuminate patterns of signal modulation by these gap junctions in future studies.

### 4) Model Limitations and Future Steps

We acknowledge several limitations in our computational modeling approach and outline the directions for future work. A major challenge lies in accurately modeling the biophysics of degeneration, as the experimental data describing deviations from healthy cellular responses remain limited. Consequently, our models rely on parameters derived from healthy cells obtained via patch clamp recordings. Incorporating biophysical data from degenerate retinas would be needed to improve predictive accuracy. Another limitation is the lack of consistency in available electrophysiological datasets. Patch clamp studies typically focus on individual cell types, and recordings of full network responses under uniform stimulation conditions are scarce. In this study, cone photocurrents were modeled using data from aquatic species, while cone bipolar cell responses were derived from rabbit and mouse. Thus, the model integrates multi-species data, which may introduce variability in neuronal responses. A unified dataset from a single species and protocol would improve model coherence. Lastly, the relatively small scale of our model and the absence of some cell types limit the true impact of photoreceptor coupling on visual coding within the network. Aberrant synapses were also observed between cell types that control the OFF pathway (GABAergic-amacrine, and OFF-cone bipolar) in RPC1 (Figure 11. Future of pathoconnectomics-simulation work will incorporate these additional cell types and synapses, while expanding the network size through learning-based approaches, to provide a more comprehensive assessment of early-stage retinal degeneration on visual signals.

**Fig 11.**
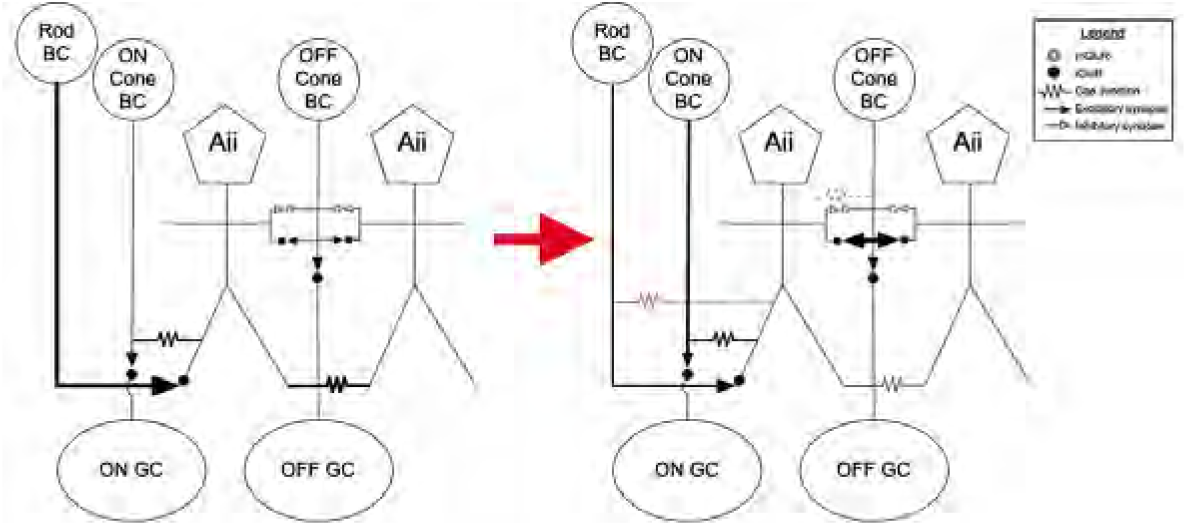
Circuit Diagram Illustrating Changes to Aii Contributions to Rod Network. Left panel is the healthy network. Right panel indicates the state of the altered network in RPC1. Weighting of lines indicates strength of input and red indicates connections only present in the degenerate retina.

Predictive computational models are valuable for developing electrotherapeutic strategies in vision research. In previous work, we used the AM-NEURON computational framework to integrate cellular with three-dimensional tissue models, enabling visualization of extracellular potentials and neuronal responses during retinal and hippocampal stimulation [34, 79, 80]. These models facilitate optimization of stimulation waveforms by establishing input-output relationships. For example, transcorneal electrical stimulation, a non-invasive therapy aimed at preserving photoreceptor integrity [81], will benefit from validated computational frameworks to predict retinal responses and optimize current delivery without extensive in vivo experimentation. Early detection of the changes in retinal activity before photoreceptor loss may provide an improved window of successful intervention and the model reported here may help facilitate the development of targeted therapeutic approaches.

## Methods and Models

### 2.4 Volume Generation

#### 2.4.1 Connectome and Pathoconnectome Volumes

The healthy connectome: Retinal Connectome 1 (RC1) first published in 2009 [82] and comprehensively described in 2011 [60] from a 13-month Dutch Belted female rabbit, contains 341 TEM sections, 18 CMP capstone (serial sections bookending the TEM volume) sections, and 11 intercalated CMP sections containing IgGs to GABA (*γ*), Glycine (G), Taurine (*τ*), Glutamate (E), and 1-amino-4-guanidobutane (AGB), which was loaded during the sample preparation. The pathoconnectome: Retinal Pathoconnectome 1 (RPC1) [35] was generated from a male 10-month transgenic P347L rabbit model of autosomal dominant RP [6, 83]. RPC1 contains 946 TEM sections with 14 intercalated CMP sections. IgGs labeled in RPC1 include GABA (*γ*), Glycine (G), Taurine (*τ*), Glutamate (E), Glutamine (Q), and Glial Fibrillary Acidic Protein (GFAP) (Table 3). After enucleation, both volumes were prepared by fixing the tissues in mixed aldehydes (1% formaldehyde and 2.5% glutaraldehyde). Tissues were then osmicated, dehydrated in graded alcohols, and embedded in epon resins. Tissues were then serially sectioned on an ultramicrotome set to 70nm and placed on formvar grids for TEM capture or on slides for CMP analysis.

**Table 3.**
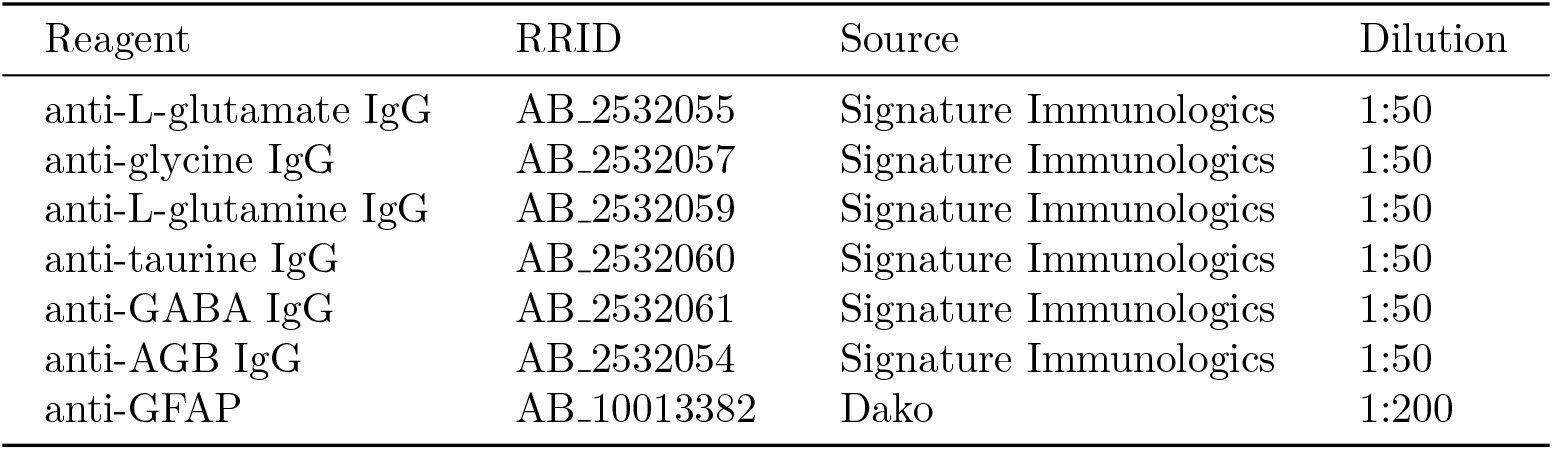
Reagents used in RPC1 and RC1 volumes.

**Table 4.** Cell IDs extracted from the RPC1 dataset and plotted in Figure 12.

| RodBCs | Aiis | CBbs | ON-RGC |
| --- | --- | --- | --- |
| 822, 933, 1001,<br>1069, 1223, 1232,<br>1242, 1243, 1258,<br>1537, 25001,<br>26167, 30692,<br>30804, 31982 | 192, 262, 265,<br>1685, 2710 | 298, 309, 430, 616,<br>617, 623, 649, 659,<br>724, 740, 1167,<br>1218, 1233, 1249 | 2612 |

#### 2.4.2 Computational Molecular Phenotyping (CMP)

Established identities of neurons in the volume are critical to understanding networks. Spatial metabolic fingerprints allow us to definitively identify cells in the volume as belonging to specific cell classes. Therefore, we include CMP analysis of small molecule and protein epitopes into each of our connectome or pathoconnectome volumes. CMP allows cell fingerprinting through the quantitative analysis of small molecule combinations, which are stoichiometrically trapped during fixation and detected with glutaraldehyde-tolerant IgGs [84–86]. The inclusion of small molecules furthers the identification of neuronal types by adding the predominant neurotransmitter as a data point to the morphology and connectivity of a given neuron. Further background on the theory of CMP can be found in prior publications [85–87].

#### 2.4.3 Image Capture, Assembly, and Annotation

Transmission electron microscopy (TEM) samples were imaged using a JEOL 1400 transmission electron microscope with a 16-Mpixel Gatan Ultrascan camera. TEM was selected because it provides high-resolution imaging (~2.18nm/px) with minimal dwell time. It also affords higher resolution recaptures with tilt for validation of individual structures such as gap junctions [36]. Samples prepared for CMP were imaged using Leica DMR light microscopes with an 8-bit CCD camera, and Surveyor software.

RC1 was assembled using the NCR toolkit [82], and RPC1 was built with a modern version of NCR Toolkit called Nornir (https://nornir.github.io/). This software assembles complete sections (mosaics) from tiled TEM or light-microscopy images and then performs semi-automated registration to align the serial sections with minimal manual correction [82, 88].

Following volume assembly, navigating the dataset and manual annotation are performed using the Viking software environment [60]. Expert annotators manually annotate every synapse (chemical or electrical) according to the parameters outlined below in the Synapse Identification and Nomenclature section. Each of these annotations is recorded in a central database which encodes each structure’s location, size, and linked partners [82]. We endeavor to minimize false positives or missed network connections through the complete annotation of all synapses. The annotation of all synapses is an audacious undertaking, and as more experts evaluate cells within a volume, missed structures may be found, leading to an evolving dataset. Both RC1 and RPC1 volumes are publicly available at

https://websvc1.connectomesdata.org/RC1/OData/ and

https://websvc1.connectomesdata.org/RPC1/OData.

### 2.5 Annotation

#### 2.5.1 Synapse Identification and Nomenclature

To detail synaptic connectivity in this study, we will refer to chemical synapses as either conventional or ribbon synapses based on their ultrastructural morphology as previously described [35]. Conventional synapses are generally inhibitory synapses of classical synaptic structure: a pre-synaptic vesicle cloud opposing a post-synaptic density. The presence of a pre-synaptic ribbon characterizes ribbon synapses, and the ribbon may oppose one or more post-synaptic densities on neighboring neurons. Ribbon synapses of the inner retina are always excitatory. Gap junctional electrical synapses are also evaluated in this study and are identified through the distinct pentalaminar structure indicative of gap junctions [35, 36]. While many gap junctions were identifiable at our native 2.18nm/pixel resolution, some were at an oblique orientation and required higher magnification recaptures with goniometric tilt. Recaptures were performed at 25,000x (0.43nm/pixel resolution).

#### 2.5.2 Morphology Annotation and Rendering

In this paper, we deviated from our classical annotations using circles [60] and used a mix of circles (in cell bodies and descending processes) and polygons to more accurately represent the morphology of the neurons presented in this study.

To render the more complex morphologies created through the addition of polygons, it was necessary to create a new morphology rendering module of Viking: VikingMesh. VikingMesh is based on the methods generated by Bajaj and colleagues for volumetric feature extraction and visualization [89]. We enhanced the results of the Bajaj methods by restricting mesh generation across sections to annotations (contours in the Bajaj terminology) known to be physically connected as recorded in the Viking database.

Contour relationships were unknown in the original Bajaj algorithm. Dae files were generated using the VikingMesh renderer (https://github.com/jamesra/Viking/tree/Legacy/MorphologyMesh), and were subsequently passed through a screened Poisson surface remeshing algorithm (https://github.com/RLPfeiffer/VikingMesh_addons) harnessing the pymeshlab python library (https://pymeshlab.readthedocs.io/en/latest/). Dae files were then imported into Blender (https://www.blender.org/) and projection images to visualize the 3D morphology and relationships were acquired, size metrics from the annotations are also preserved throughout the rendering process.

### 2.6 Data Extraction and Simulation Framework

The NEURON simulation environment was used generate a retinal network from the annotated pathoconnectome volume. Neuronal responses to light stimulation were simulated by computing membrane potentials at each time step for a specified current-clamp input. The model combines the baseline activity of a healthy retina with the synaptic rewiring observed in RPC1 to simulate the effects of degeneration on network-level signaling. Modified versions of membrane models from other studies presenting single-compartment and multi-compartmental retinal cells were used to generate the baseline model. A light input stimulation paradigm that simulates the natural photocurrent generated after a light flash is also introduced.

#### 2.6.1 Morphology and Topology

For our modeling paradigms, all cells, except photoreceptors, were modeled multi-compartmentally to accurately represent the soma, axon, and dendritic branches. Fourteen RodBCs, thirteen CBbs, five Aiis and one RGC were extracted along with their synaptic mappings from the RPC1 dataset to build the network model (Table 4). The number of cell extracted was in part influenced by the completeness of the cell within the volume; cells with large processes extending off volume were excluded from the analysis. The 3-D representation of the network is shown in Figure 12, plotted using VikingMesh.

**Fig 12.**
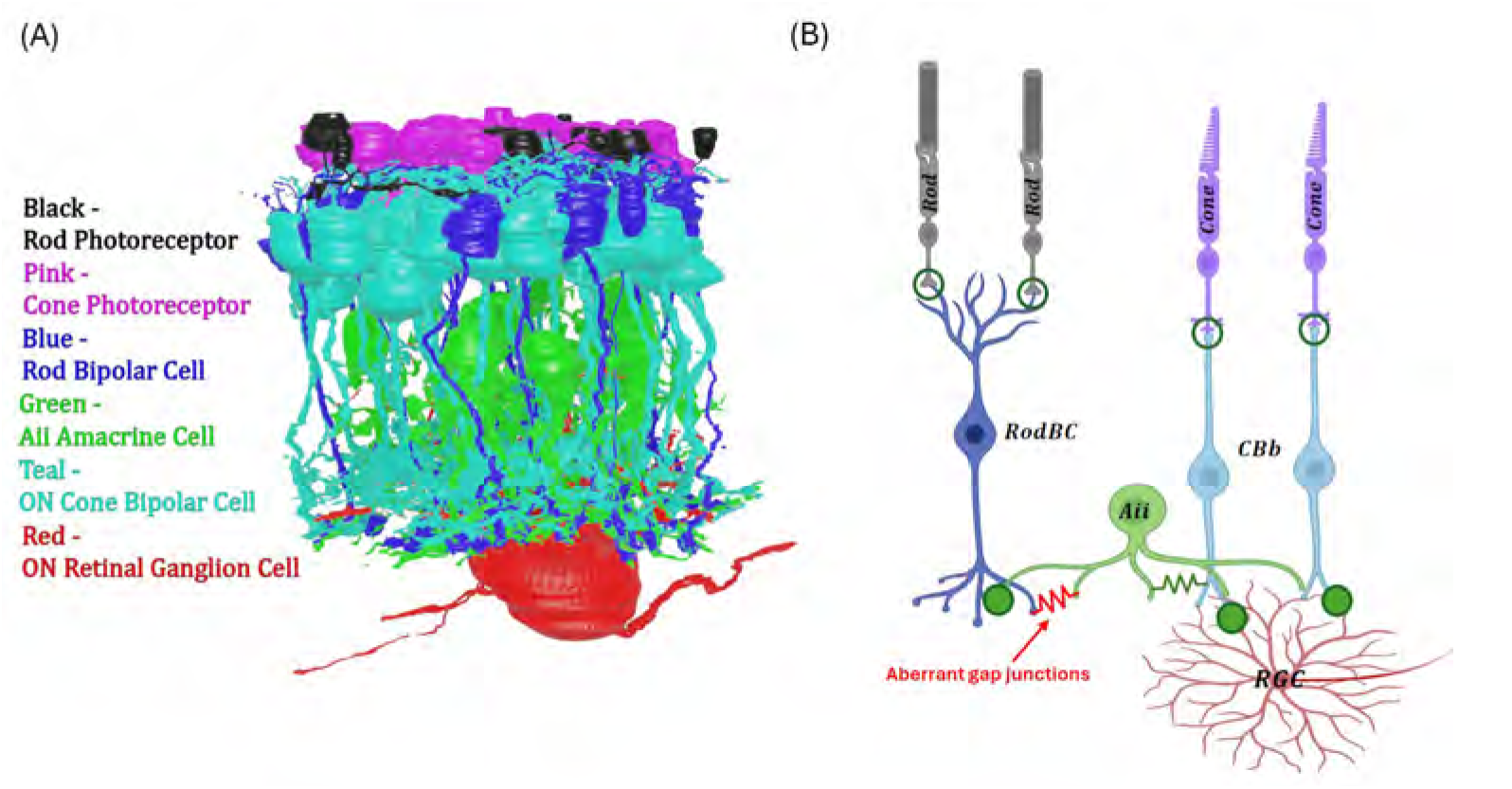
(A) 3D plot of the RPC1 cell network, showing the morphology and topology captured by TEM imaging. (B) Signal flow illustration in the ON-type RPC1 network. Green connections are normally found in a healthy retina, while red connections appeared only in the pathoconnectome. Empty circles represent ribbon synapses with metabotropic mGluR6 channels, solid circles represent ionotropic iGluR channels, and resistors represent gap junctions.

#### 2.6.2 Pre-processing and Simulation in NEURON

A Python script was used to download cell morphologies from the connectome database in .swc format and their direct (one-hop) synaptic connections in .txt format. The .swc files were processed in MATLAB to ensure a single root, bridge missing edges, fix annotations, and correctly assign compartment types (soma, axon, dendrite). The processed .swc files and synaptic data were then used to create the network model. Membrane potential responses to photocurrent (current-clamp) stimulation were solved in NEURON for each cell type. The compartmentalization system followed cable theory principles [90]. Each compartment was defined as a tapered cylinder connected to neighboring compartments by an intracellular resistor and assigned passive or active membrane currents represented by equivalent electrical circuits (Figure 13). Network topology and synaptic mapping were downloaded along with the morphology of the selected cells. Synapses were defined by type (ribbon or gap junction) and weight (the post-synaptic or gap junction density area).

**Fig 13.**
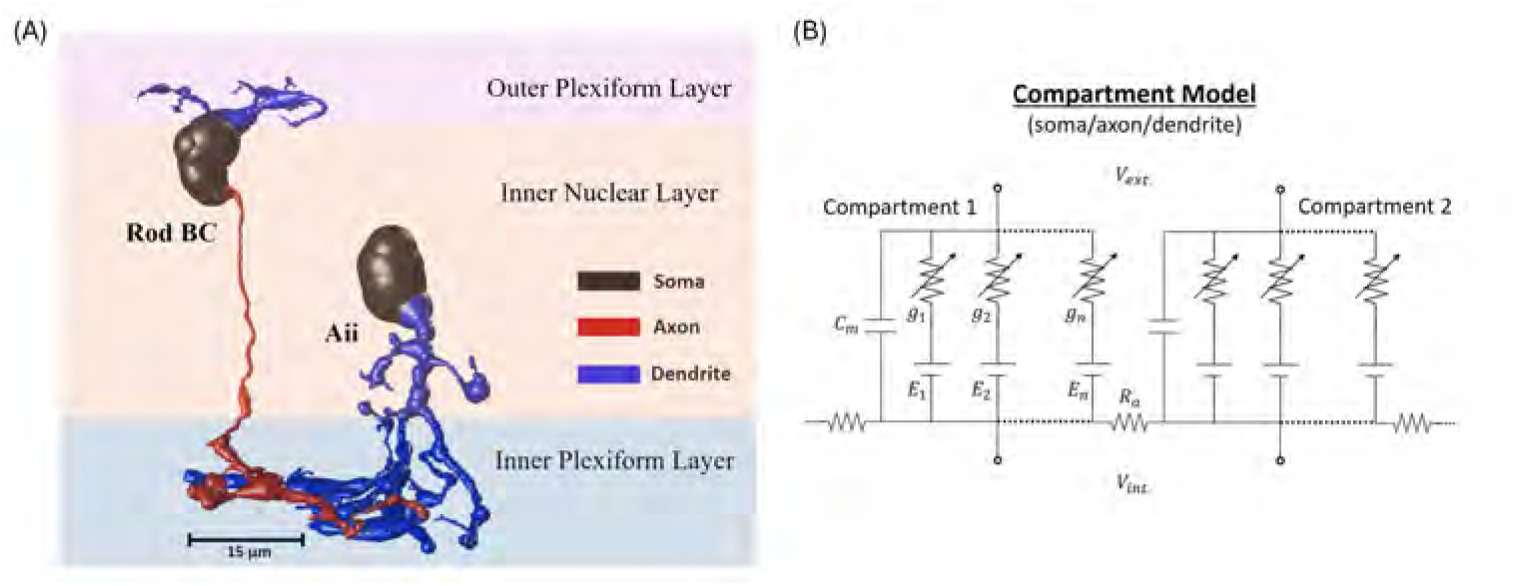
Multi-compartment cable model of retinal neurons. (A) Examples of RodBC (# 822) and Aii (# 2710) morphologies from RPC1 are shown in detail. The cells are mapped in 3-D, corresponding to their realistic positions in the retina. (B) Each compartment is defined by type (soma/axon/dendrite) with assigned membrane conductance and capacitance. Adjacent compartments are connected through axial resistance. The shift in membrane potential (*V*_*ext*_ −*V*_*int*_) modulates the current flow across membrane channels.

#### 2.6.3 Light Stimulation Protocol

The stimulation protocol is based on a method developed to reproduce natural light responses in the cone pathway [91]. This method was validated using patch-clamp recordings of photoreceptor photocurrents [92–95], as well as corresponding membrane potential changes in photoreceptors [96, 97] and bipolar cells [98–100]. Photoreceptors were modeled as point sources with static photocurrent waveforms applied through a current clamp at the inner segment. These waveforms capture the current generated at the inner segment during natural phototransduction at the outer segment.

This stimulation protocol was optimized to match the recorded EPSPs of RodBCs, CBbs, Aiis, and RGCs to their light-evoked responses [97, 98, 101, 102], while maintaining network convergence ratios [103, 104]. Cones exhibit a faster photocurrent decay, repolarizing within 0.5 s after a light flash, compared to 5 s for rods. Their photocurrent amplitude is also smaller, saturating at 20 pA, compared to 30 pA in rods [92, 105, 106].

Additionally, a noise model was added to the photocurrent waveform, to reproduce the spontaneous activity in dark-adapted retina. The model consists of a nonlinear stimulus-evoked current with a sigmoidal onset and an adjustable plateau duration, combined with continuous stochastic fluctuations. Stochastic variability is introduced multiplicatively, allowing fluctuations to scale naturally with the operating level of the current. As a result, both dark-adapted current and light-evoked responses exhibit smooth, physiologically realistic variability without introducing discontinuities at stimulus transitions.

#### 2.6.4 Current Injection Model

The injected current *I*(*t*) consists of a baseline component with stochastic fluctuations and a time-dependent stimulus component applied during discrete activation periods.

#### Baseline current and noise

In the absence of stimulation, the injected current is given by

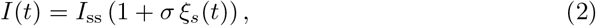

where *I*_ss_ is the steady-state baseline current, *σ* is the noise amplitude, and *ξ*_*s*_(*t*) is a temporally correlated stochastic process. The noise is modeled as an Ornstein–Uhlenbeck (OU) process:

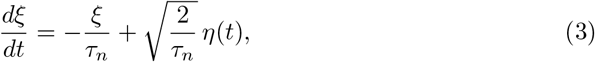

where *τ*_*n*_ is the noise correlation time constant and *η*(*t*) is Gaussian white noise with zero mean and unit variance. The OU process is subsequently low-pass filtered:

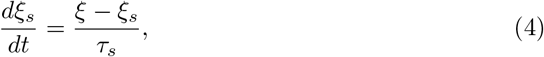

where *τ*_*s*_ is a smoothing time constant. The first-order low-pass filter is applied to ensure the noise does not have sharp edges, which could cause an out of bounds error. Without the filter, the simulation requires higher temporal resolution, significantly increasing the computation time.

#### Stimulus current

During stimulation, the injected current is defined as

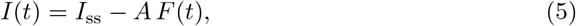

where *A* is the stimulus amplitude and *F* (*t*) is a nonlinear time-dependent waveform. Let *t*_*r*_ denote the time since stimulus onset:

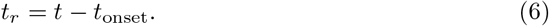

The waveform is composed of four components:

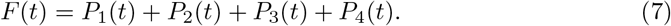

#### Waveform shape components

A sigmoidal activation function is defined as

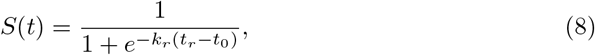

and the primary rising component is given by

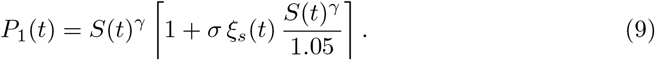

The remaining components shape the temporal profile of the stimulus:

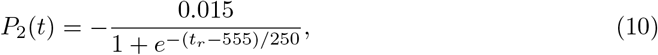

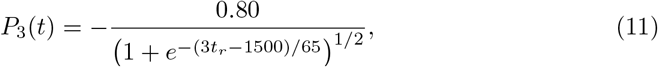

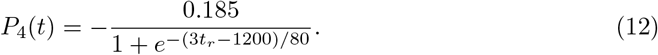

#### Temporal structure

Stimulation is delivered as a sequence of *N* pulses. Each pulse has duration *T*_on_ and is followed by an inter-pulse interval *T*_off_, with an initial delay *T*_delay_ before the first pulse.

Thus, the injected current alternates between:

- an *active state*, where *I*(*t*) = *I*_ss_ −*AF* (*t*),
- an *inactive state*, where *I*(*t*) = *I*_ss_ (1 + *σξ*_*s*_(*t*)).

The stochastic dynamics are implemented using an explicit time-stepping scheme, and therefore assume a fixed integration time step. The resulting noise spectrum exhibits enhanced attenuation at high frequencies while remaining stationary and temporally correlated. The OU process was selected because it provides a minimal biophysically interpretable model of the intrinsic noise of retinal circuits using a single correlation parameter.

### 2.7 Cell Biophysical Models

Membrane properties for each cell type were defined using published models in the ModelDB database. Each cell was modeled as multi-compartmental, with cell type–specific membrane properties assigned to each compartment (Figure 13). Voltage-gated ion channels were modeled by conductance-based kinetic models derived from patch-clamp recordings of individual cells. Because experimental data describing degenerate cell membrane properties is limited, the biophysical parameters used here are from healthy cells. Although studies of early-stage RP have shown reduced electroretinogram amplitudes in photoreceptors and bipolar cells [107, 108], ERG lacks the precision of patch-clamp data for computational modeling of membrane responses. Therefore, these findings were not incorporated in the available models.

#### 2.7.1 Cell-specific Parameters

Whenever possible, the numbers and relative strengths of the cells and synapses were obtained from the RPC1 dataset. However, the uncertainty of the identity of a subset of photoreceptor terminals in RPC1 described in the initial RPC1 manuscript [35], necessitated the approximation of photoreceptor inputs used in this model. The parameters used for each of the respective cell classes are described in Supporting Information S1.

### 2.8 Synaptic Models

The network model of this study includes only the ON pathway (Figure 12), which comprises metabotropic and ionotropic ribbon synapses, as well as electrical synapses mediated by gap junctions. Ribbon synapses were modeled as either graded, occurring at the postsynaptic terminals of bipolar cells with photoreceptors and at the postsynaptic terminals of RGCs [109], or as exponential, capturing the fast, transient potential changes of Aiis [110]. Gap junctions, the anatomical basis of electrical synapses between neurons and bidirectional by nature in vertebrates [111], were used to establish electrical coupling. Because not all cells are fully contained within the imaged volume, the computational model has some discrepancy from the full synaptic information presented in section 1.3.5. Namely, RodBCs 31982, 45599, 41743 and 22748 were not included, and their gap junctions with Aiis had to be omitted. Thus, the network model has 37 RodBC::Aii gap junctions, as opposed to the full 44 imaged in the volume. The count of each synapse of the ON pathway can be found in Table S2.

Individual values for each biophysical parameter such as membrane channel conductances and synaptic thresholds are provided in the open-access GitHub repository. The RPC1 model files with the complete parameters and scripts can be accessed and simulated using the instructions in the following link:

https://github.com/LazziLab/Pathoconnectome-Modeling

## Acknowledgments

The work has been supported by the NIH grants R01 EY035527, R01 EY028927, P30 EY08098, and an unrestricted grant from Gabe Newell. BWJ is a recipient of an Research to Prevent Blindness Stein Innovation Award, and an Unrestricted Research Grant from Research to Prevent Blindness, New York, NY to the Department of Ophthalmology University of Pittsburgh. Additionally, RLP is a recipient of the Research to Prevent Blindness Career Development Award.

