## SupplementalMethods for "Impact of Aii Amacrine Cell Rewiring in a Pathoconnectome-Based Computational Model of Early Retinal Degeneration"

### Supporting Information

#### S1 Biophysical Cell Parameters.

##### Rod Photoreceptor

Reports on the number of rod inputs per rod bipolar cell (RodBC) in the healthy rabbit retina vary widely. One study suggests that up to thirty rods provide input to a single RodBC [1], whereas another shows that approximately fifty rods fall within the dendritic field of a RodBC, with each rod potentially contacting multiple RodBCs [2]. Other studies estimate this number to be closer to one hundred on average [3, 4].

In our previous computational study evaluating the membrane depolarization of morphologically altered RodBCs [5], we used thirty rods per cell, based primarily on mouse circuitry data [6, 7]. Consistent with RP pathology, in which rod photoreceptors are the first to undergo stress and apoptosis [8], pathoconnectomics data revealed a diminished rod count. Adjacent vertical sections indicate approximately 50% rod loss in the outer nuclear layer [9]. Based on this reduction, along with the inconclusive identification of certain photoreceptor inputs, we infer that rod signal strength may be reduced by up to 50%. Assigning an average of fifteen rods to each RodBC is therefore statistically plausible at this stage of degeneration and enables systematic evaluation of how novel RodBC connectivity affects signaling in the inner plexiform layer. Additional simulations with varying numbers of rod inputs per RodBC were performed to assess how input count influences RGC output; these results are discussed in section 2.3. Model precision can be further improved as more comprehensive annotations become available. The rod model was adapted from an ON-type retinal network [10]. A transient light stimulation protocol, described in our previous work [5, 11], was used to define the photocurrent waveform. This approach, commonly used in patch-clamp experiments, replicates currents measured in rod photoreceptors in response to a 10 ms, 500 nm light flash [12, 13].

Six membrane currents were used to model changes in rod membrane potential at the inner segment in response to the photocurrent: hyperpolarization-activated current ($I_h$), non-inactivating potassium current ($I_{Kx}$), delayed rectifier potassium current ($I_{Kv}$), calcium current ($I_{Ca}$), calcium-dependent chloride current ($I_{Cl(Ca)}$), and calcium-dependent potassium current ($I_{K(Ca)}$). Together, these currents reproduce the membrane hyperpolarization observed during photocurrent activation (Fig. 9B). Compartment dimensions were tuned to reproduce experimentally observed membrane potential responses. Because photoreceptors were modeled as point sources, compartment size can be adjusted to achieve optimal agreement with experimental data. The rod compartment was set to have a radius of 1 $\mu$m and a length of 1 $\mu$m to match the target membrane potential response. A MATLAB script was used to randomize the spatial distribution of photoreceptors and establish synaptic connections across RodBC dendritic terminals.

##### Cone Photoreceptor

Membrane currents present in rods also exist in cones, except $I_{Kv}$ [14]. Differences in channel conductance and reversal potentials result in a weaker hyperpolarization response in cones (Fig.9B). As with rods, the cone compartment size is tuned to reproduce the target membrane potential response from patch-clamp recordings [11], set to a radius of 2.25 $\mu$m and a length of 2 $\mu$m. Since cone synapses with cone bipolar cells (CBbs) are not yet completed in the RPC1 dataset, literature estimates were used to determine the average number of cone inputs per bipolar cell [6,15]. Based on these studies, each CBb was connected to eight cones—an upper estimate appropriate for the peri-streak region represented in the RPC1 volume, where cone density is higher. Synaptic locations were randomized using a MATLAB script, like with the rod inputs.

#### Rod Bipolar Cell

RodBCs were modeled using the following ionic currents: voltage-dependent potassium ($I_{Kv}$), calcium-dependent potassium ($I_{K(Ca)}$), calcium ($I_{Ca}$), hyperpolarization-activated ($I_h$), and transient outward ($I_A$) currents [10,16,17]. These were implemented using modified Hodgkin–Huxley equations, with kinetic parameters tuned to reproduce electrophysiological recordings from rat RodBCs [18–20] and voltage-clamp data [21].

#### ON-Cone Bipolar Cell

CBbs form the primary visual pathway by directly connecting to RGCs and receiving input from cone photoreceptors. Multiple subtypes were identified in rat [22], mouse [23–25], and human [26], some of which correspond to rabbit classes [27]. While the pathoconnectome data allow differentiation between RodBCs and CBbs, incomplete class annotation among CBb subtypes prevents a more comprehensive classification. Therefore, we adopted a general model for the depolarizing CBb, which is non-spiking [10]. This model shares the same membrane channels as RodBCs, excluding $I_A$ (S1 Table).

#### Aii Amacrine Cell

The Aii is a key component in the rod pathway that couples rod signals with with CBbs through gap junctions. Current-clamp data suggest that Aiis exhibit two distinct components in their membrane potential response: a transient, large-amplitude depolarization immediately after stimulation, followed by a sustained depolarization of lower amplitude that coincides with the duration of CBb depolarization shared through bidirectional gap junctions [28–32]. Aiis express Hodgkin–Huxley sodium ($I_{HHNa}$) and potassium ($I_{HHK}$) channels, as well as calcium ($I_{Ca}$) and A-type potassium ($I_{Ka}$) channels [10,33]. Passive membrane parameters were modified to match the curve-fitted, electrically coupled Aii model [34].

#### ON-Retinal Ganglion Cell

RGCs are the most extensively studied retinal neurons, responsible for visual output and transmitting it to the visual cortex via the optic nerve. Our model is constructed around a single RGC that integrates the signals originating from the photoreceptors, which get processed through the network. The RGC responds to light stimulation by generating a train of action potentials, with firing rate dependent on the photocurrent amplitude [35].

The RGC membrane kinetics are based on a model from [36], previously used to study targeted electrical stimulation of RGC subtypes [37]. This biophysical model does not account for spontaneous firing observed in the absence of stimulation [38]; therefore, a noise component was added to the input current to simulate the resting activity, described in section 2.6.4.

**S1 Table. Biophysical Membrane Properties for Retinal Cells**

| Cell Type | Ionic Channels | Reference Model |
|---|---|---|
| Rod Photoreceptor | $I_h$,$I_{Kx}$,$I_{Kv}$,$I_{Ca}$,$I_{Cl(Ca)}$,$I_{K(Ca)}$ | [10, 17] |
| Cone Photoreceptor | $I_h$,$I_{Kv}$,$I_{Ca}$,$I_{Cl(Ca)}$,$I_{K(Ca)}$ | [11, 14] |
| Rod Bipolar | $I_h$,$I_{Kv}$,$I_{Ca}$,$I_A$,$I_{K(Ca)}$,$I_{leak}$ | [10, 16] |
| Cone Bipolar | $I_h$,$I_{Kv}$,$I_{Ca}$,$I_A$,$I_{K(Ca)}$,$I_{leak}$ | [10, 11] |
| Aii Amacrine | $I_{HHNa}$,$I_{HHK}$,$I_{Ca}$,$I_{Ka}$ | [10, 33, 34] |
| Retinal Ganglion | $I_h$,$I_{Kv}$,$I_{Ca}$,$I_{K(Ca)}$,$I_{leak}$ | [36] |

### S2 Modeled Synaptic Properties

#### Rod Photoreceptor – Rod Bipolar Cell

We used the ribbon synapse model presented in Publio's work [17], a simplified version of Mulloney's graded synapse model [39], where the parameters were adapted from Sikora's ribbon synapse model [40]. This glutamatergic synapse depends on the rate of glutamate release by photoreceptors, producing a graded voltage response in bipolar cells. Channel conductances from the reference models were tuned to better match the EPSP shape observed in patch-clamp recordings.

#### Cone Photoreceptor – ON Cone Bipolar Cell

Like the rod photoreceptor, the cone synapse is driven by glutamate release and modeled as a graded response. The same synaptic model used for the rod-RodBC connection was applied, with the synaptic weight adjusted to produce realistic EPSPs in CBbs consistent with experimental recordings [11].

#### Cone Photoreceptor – Rod Photoreceptor

Although some published network models have included electrical coupling between rods and cones [10, 41], we excluded it here because the imaged volume does not extend beyond the base of the outer nuclear layer, preventing synaptic mapping as done for other cell types. Additionally, since photoreceptors were randomly distributed along bipolar cell terminals in our model, the density of rod–cone coupling could not be reliably estimated as reported in the literature [42]. Experimental evidence indicates that connexin 36 is the dominant protein mediating these bidirectional connections [43, 44].

#### ON Cone Bipolar Cell – Aii Amacrine Cell

Aiis are postsynaptic to ribbon synapses from RodBCs and electrically coupled to CBbs via GJs, as confirmed by the connectomics data. Gap junctions form direct electrical couplings between cells, and their blockade has been observed to reduce the membrane potential time constant [45, 46], suggesting a linear resistive relationship for the equivalent circuit model. Thus, GJs were modeled as resistors with a conductance of 200 pS, consistent with previous modeling work [10, 47]. Some rodent physiology studies suggest higher average conductance values around 700 pS [48, 49], while more recent evidence indicates values closer to 400 pS in rat [50] or 500 pS in mouse [51]. We used

200 pS across all GJs in our core simulations for consistency with prior models; the effect of higher conductance values is examined in section 2.3.2.

#### Aii Amacrine Cell – Aii Amacrine Cell

Aiis in the network are also electrically coupled via GJs, which may enhance the signal-to-noise ratio within the rod pathway by amplifying correlated rod signals and suppressing uncorrelated noise [44]. The extent of Aii coupling varies with light adaptation, from about 20 cells in dark-adapted to over 300 in light-adapted rabbit retina [31]. Tight coupling among a few cells enables single-photon signal detection by filtering inactive regions during scotopic conditions, thereby increasing sensitivity [45,52]. However, Aiis uncouple following intense background illumination, possibly due to increased D1 receptor activation [31]. The present model includes only five coupled Aiis, making it best suited for simulating dark-adapted or light-saturated conditions.

#### ON Cone Bipolar Cell – Ganglion Cell

CBbs link the cone and rod pathways via Aiis. The CBb is the only presynaptic input to the RGC and modulates its firing rate. The synapse is graded with first-order kinetics, similar to the photoreceptor–bipolar cell synapses, with equilibrium potential and conductances adjusted to match light-evoked RGC responses [36]. Due to missing annotations between these two cell types, we adopted a strategy similar to that used for the photoreceptors, creating artificial synapses based on annotations from a healthy rabbit connectome dataset [53]. We averaged the number and weights of CBb-RGC synapses and incorporated them into our topology, assuming minimal rewiring within the ganglion cell layer during early degeneration.

#### Rod Bipolar Cell – Aii Amacrine Cell

In the healthy retina, signal flow from RodBCs to Aiis occurs unidirectionally through excitatory ribbon synapses [53] and the Aiis' arboreal dendrites are always postsynaptic to RodBCs [54]. This synaptic connection was modeled using ionotropic glutamate receptors (iGluR), exhibiting fast depolarization and slower repolarization upon excitation, as shown by light-response recordings [28–31,33]. NEURON's exponential synapse function `ExpSyn` was used to model these iGluR-mediated connections. During early-stage degeneration, RodBC–Aii connections undergo rewiring and form aberrant GJs not observed in wild-type rabbit [9]. In the absence of molecular or physiological data, these aberrant junctions were modeled identically those of Aii–CBb, as passive resistive elements with a conductance of 200 pS.

#### Missing Cell Types

Another amacrine cell type in RPC1, a subset of GABAergic amacrine cells, exhibits rewiring during early degeneration, in part, by extending aberrant processes into the outer plexiform layer [55]. Due to the lack of published models and patch-clamp data for this cell type, it was not included in the present network.

**S2 Table. Synaptic distribution in the network model between neighboring cells. ($R_g$): metabotropic graded ribbon, ($-R_g$): ionotropic graded ribbon, ($R_e$): excitatory exponential ribbon, (G): gap junction.**

| 1-Hop Connections | Baseline Synapses | New Synapses due to Rewiring |
|---|---|---|
| Rods - RodBCs | 210 ($R_g$) | 0 |
| Cones - CBbs | 104 ($R_g$) | 0 |
| RodBCs - Aiis | 197 ($R_e$) | 37 (G) |
| Aiis - Aiis | 71 (G) | 0 |
| Aiis - CBbs | 80 (G) | 0 |
| CBbs - RGC | 13 ($R_g$) | 0 |
